# Influence of species and processing parameters on recovery and content of brain tissue-derived extracellular vesicles

**DOI:** 10.1101/2020.02.10.940999

**Authors:** Yiyao Huang, Lesley Cheng, Andrey Turchinovich, Vasiliki Mahairaki, Juan C. Troncoso, Olga Pletniková, Norman J. Haughey, Laura J. Vella, Andrew F. Hill, Lei Zheng, Kenneth W. Witwer

**Affiliations:** Department of Molecular and Comparative Pathobiology, Johns Hopkins University School of Medicine, Baltimore, Maryland, USA; Department of Laboratory Medicine, Nanfang Hospital, Southern Medical University, Guangzhou, Guangdong, China; Department of Biochemistry and Genetics, La Trobe Institute for Molecular Science, La Trobe University, Bundoora, Australia; Molecular Epidemiology, German Cancer Research Center DKFZ, Heidelberg, Germany; SciBerg e.Kfm, Mannheim, Germany; Department of Neurology, Johns Hopkins University School of Medicine, Baltimore, Maryland, USA; Department of Pathology, Johns Hopkins University School of Medicine, Baltimore, Maryland, USA; The Florey Institute of Neuroscience and Mental Health, The University of Melbourne, Parkville, Australia; Department of Surgery, The University of Melbourne, The Royal Melbourne Hospital, Parkville, Victoria 3050, Australia

**Keywords:** Extracellular vesicles, brain, central nervous system, tissue preparation, postmortem interval, small RNA sequencing, proteomics, exosomes, neurodegenerative disease

## Abstract

Extracellular vesicles (EVs) are involved in a wide range of physiological and pathological processes by shuttling material out of and between cells. Tissue EVs may thus lend insights into disease mechanisms and also betray disease when released into easily accessed biological fluids. Since brain-derived EVs (bdEVs) and their cargo may serve as biomarkers of neurodegenerative diseases, we evaluated modifications to a published, rigorous protocol for separation of EVs from brain tissue and studied effects of processing variables on quantitative and qualitative outcomes. To this end, size exclusion chromatography (SEC) and sucrose density gradient ultracentrifugation were compared as final separation steps in protocols involving stepped ultracentrifugation. bdEVs were separated from brain tissues of human, macaque, and mouse. Effects of tissue perfusion and a model of post-mortem interval (PMI) before final bdEV separation were probed. MISEV2018-compliant EV characterization was performed, and both small RNA and protein profiling were done. We conclude that the modified, SEC-employing protocol achieves EV separation efficiency roughly similar to a protocol using gradient density ultracentrifugation, while decreasing operator time and, potentially, variability. The protocol appears to yield bdEVs of higher purity for human tissues compared with those of macaque and, especially, mouse, suggesting opportunities for optimization. Where possible, perfusion should be performed in animal models. The interval between death/tissue storage/processing and final bdEV separation can also affect bdEV populations and composition and should thus be recorded for rigorous reporting. Finally, different populations of EVs obtained through the modified method reported herein display characteristic RNA and protein content that hint at biomarker potential. To conclude, this study finds that the automatable and increasingly employed technique of SEC can be applied to tissue EV separation, and also reveals more about the importance of species-specific and technical considerations when working with tissue EVs. These results are expected to enhance the use of bdEVs in revealing and understanding brain disease.

## Introduction

Extracellular vesicles (EVs) are nano-sized, lipid bilayer-delimited particles that are released by various cells. They can package and deliver molecules such as RNAs and proteins and are thus involved in multiple physiological and pathological pathways by serving as messengers in cell-to-cell communication. Roles of EVs in the central nervous system (CNS) have now been well established. EVs are released by all neural cells^1–3^, including neurons, oligodendrocytes, astrocytes, and microglia. They can carry disease-associated agents such as amyloid-beta (Aβ)^4–7^ and tau^6–8^ proteins, which may promote neurodegenerative and inflammatory diseases. However, EVs may also exert protective functions in the CNS by distributing anti-inflammatory factors^9–11^. While brain-derived EVs (bdEVs) may leave the brain and betray the state of the CNS as biomarkers in blood and other peripheral fluids, bdEVs are first found in the tissue interstitial space^12^ and may be most likely to act locally. The composition of tissue EVs may thus shed light on physiological and pathological mechanisms in the brain. Moreover, bdEVs in tissue could be used to identify reliable cell-specific markers that could then be used to capture specific populations of CNS-origin EVs in the periphery, helping to diagnose and monitor CNS disease.

In separating EVs from post-mortem brain tissue, as from any tissue, it is critical to achieve some degree of tissue disruption while minimizing cellular disruption; to impose one or more EV separation steps to increase purity; and to show that EVs have been enriched with minimal cellular (or other) contamination. If cells are destroyed, intracellular components and artificially produced vesicles may co-isolate with EVs^13^. Gentle tissue separation might include mechanical (slicing), enzymatic digestions^12–16^, and/or immersion in cell culture medium ^17–19^. However, extensive mincing or grinding/homogenization, as reported in several studies ^12, 15, 20–22^, may result in non-EV contaminants and challenge the definition of tissue EVs. Following tissue preparation EV separation is done. Reported methods include standard ultracentrifugation (UC)^21^, density gradient ultracentrifugation (DGUC)^12, 13, 15^, chemical separation/precipitation^19, 20, 23^ or combinations of these techniques. Finally, after separation, thorough characterization of both EVs and potential contaminants is needed, but only some groups consider the latter ^13–15, 22^.

Previously, a DGUC method with a triple sucrose cushion was used as a final step in rigorous separation of EVs from brain tissue^13^. By careful characterization of enriched and depleted EV components, the published method was shown to be highly effective at eliminating cellular contaminants that do not have the same density as EVs. The method is thus valuable and effective, but the DGUC step can also be relatively time-consuming^24–26^ (at least three hours), requires an operator skilled in preparing density gradients, and for this reason may have some between-run or between-operator variation. We thus queried if size exclusion chromatography (SEC), an automatable and increasingly employed method of EV separation, could be used in place of DGUC to achieve acceptable EV purity following filtration and 10,000 x g centrifugation. In this study, we applied an SEC-containing protocol to separate small EVs (sEVs) from brain tissues of human, macaque, and mouse. We also evaluated the effects of tissue perfusion and post-mortem interval before final bdEV extraction on various parameters of the recovered bdEVs. Small RNA sequencing and proteomics revealed molecular profiles of bdEVs of human and macaque. In summary, this study adds size exclusion to the list of techniques that can be applied in bdEV separation and evaluates numerous factors affecting bdEV separation from brain of different species.

## Methods

### Tissue Collection and Preparation

Post-mortem tissues were obtained from separate studies: human (Table 1, Parietal cortex, frozen, Johns Hopkins Alzheimer’s Disease Research Center), mouse (Table 2, C57BL/6 mice, whole brains without cerebellum and olfactory bulbs), fresh, perfused and non-perfused), and macaque (Table 3, occipital lobe, perfused, fresh, different post-mortem interval). All procedures were approved by the Johns Hopkins Univeristy Institutional Review Board (human samples) or the Johns Hopkins University Institutional Animal Care and Use Committee (IACUC, animal studies) and conducted in accordance with the Weatherall Report, the Guide for the Care and Use of Laboratory Animals, and the USDA Animal Welfare Act. For human tissue, following external examination and weighing of the autopsy brain, the right cerebral hemisphere was cut into coronal slabs, frozen on pre-chilled metal plates, and stored at −80°C. For macaque, fresh occipital lobes were divided into three pieces and stored in Hibernate-E medium (Thermo Fisher A12476-01) for 2 hours, 6 hours, and 24 hours, respectively, at room temperature before EV separation to investigate effects of post-mortem interval (PMI) ^27^. For mouse, whole brains with or without perfusion with PBS (Thermo Fisher 14190250) through the left ventricle were collected. A total of 50 ml PBS was used for perfusion at a speed of 20 ml/min. Perfused and non-perfused mouse brains were snap frozen on dry ice. All tissues were stored at −80°C before use unless otherwise noted.

**Table 1.**
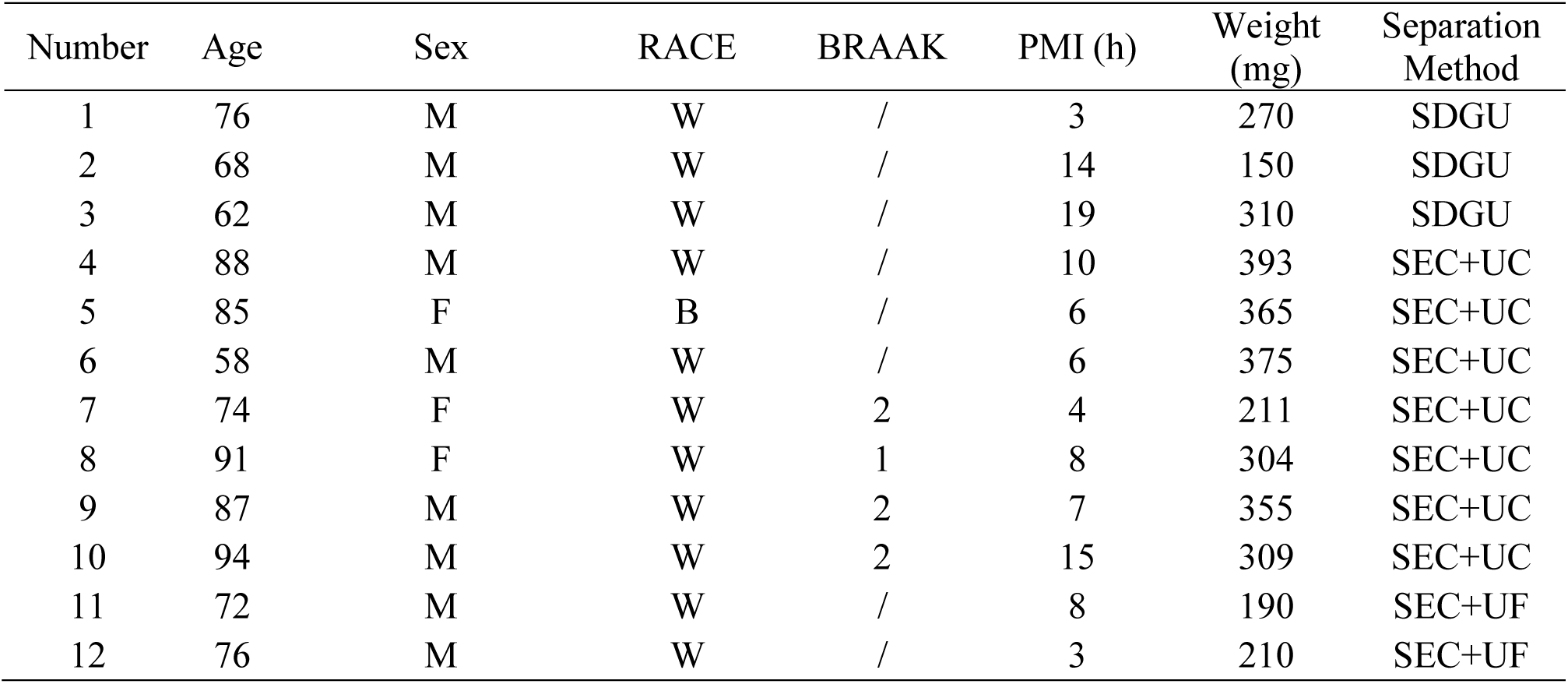
Human cortical samples

| Number | Age | Sex | RACE | BRAAK | PMI (h) | Weight (mg) | Separation Method |
| --- | --- | --- | --- | --- | --- | --- | --- |
| 1 | 76 | M | W | / | 3 | 270 | SDGU |
| 2 | 68 | M | W | / | 14 | 150 | SDGU |
| 3 | 62 | M | W | / | 19 | 310 | SDGU |
| 4 | 88 | M | W | / | 10 | 393 | SEC+UC |
| 5 | 85 | F | B | / | 6 | 365 | SEC+UC |
| 6 | 58 | M | W | / | 6 | 375 | SEC+UC |
| 7 | 74 | F | W | 2 | 4 | 211 | SEC+UC |
| 8 | 91 | F | W | 1 | 8 | 304 | SEC+UC |
| 9 | 87 | M | W | 2 | 7 | 355 | SEC+UC |
| 10 | 94 | M | W | 2 | 15 | 309 | SEC+UC |
| 11 | 72 | M | W | / | 8 | 190 | SEC+UF |
| 12 | 76 | M | W | / | 3 | 210 | SEC+UF |

**Table 2.**
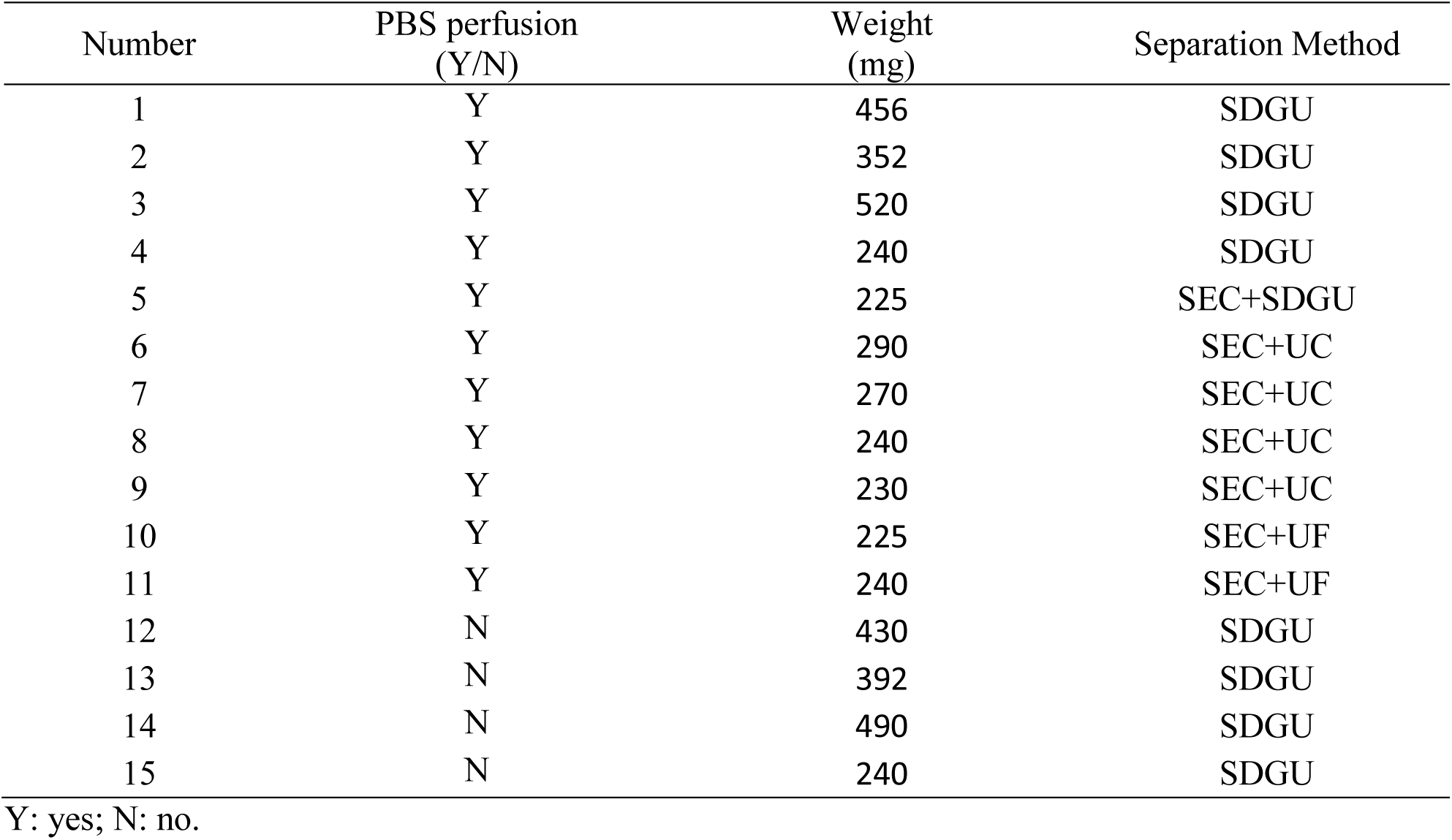
Mouse brain samples

**Table 3.** *M. nemestrina* occipital lobe samples

| Subject Number | Sex | SIV Status | Days post inoculation | Post-mortem Interval (PMI, h) | Weight (mg, by PMI) | Separation Method |
| --- | --- | --- | --- | --- | --- | --- |
| 1 | M | Infected | 49 | 2, 6, 24 | 480, 450, 440 | SEC+UC |
| 2 | M | Infected | 51 | 2, 6, 24 | 460, 400, 430 | SEC+UC |
| 3 | M | Infected | 44 | 2, 6, 24 | 504, 480, 507 | SEC+UC |
| 4 | M | Infected | 46 | 2, 6, 24 | 450, 443, 458 | SEC+UC |

Sex: M = male; no samples from females

### Separation of Extracellular Vesicles from Tissue

EVs were separated from tissue using the protocol established previously by Vella, et al, ^13^ with minor modifications. Before extraction, a small (∼50 mg) piece of tissue was stored at −80°C for later protein and RNA extraction. The remaining frozen tissue was weighed and briefly sliced on dry ice and then incubated in 75 U/ml collagenase type 3 (Worthington #CLS-3, S8P18814) in Hibernate-E solution (human tissue for 20 min, macaque and mouse tissue 15 min, based on previous optimizations of the Vella and Hill laboratories and our findings that macaque and mouse brains are more fragile and sensitive to digestion compared with human tissue) at 37°C. PhosSTOP and Complete Protease Inhibitor (Sigma-Aldrich PS/PI 4906837001/11697498001) solution was then added to stop digestion. The dissociated tissue was spun at 300 × g for 10 min at 4°C. Small pieces of the pellet (“brain homogenate with collagenase, BHC”) were stored at −80°C for later protein extraction, while supernatant was transferred to a fresh tube and spun at 2000 × g for 15 min at 4°C. Cell free supernatant was filtered through a 0.22 μm filter gently and slowly (5 ml of supernatant passed through the filter over approximately 1 min) (Millipore Sigma, SLGS033SS) for further depletion of cell debris and spun at 10,000 × g for 30 min at 4°C (AH-650 rotor, Beckman Ultra-Clear Tube with 5 ml capacity). The pellet was resuspended in 100 μl PBS and considered to be the “10K pellet” (10K or 10K fraction). The 10,000 × g supernatant was then processed by sucrose density gradient ultracentrifugation (SDGU), as previously described ^13^, or by SEC followed by concentration by ultracentrifugation (UC) or ultrafiltration (UF).

### Sucrose Density Gradient Ultracentrifugation (SDGU)

For SDGU, the sEV-containing supernatant from the 10,000 x g step was overlaid on a triple sucrose cushion (F2: 0.6 M, 1.0810 g/cm^3^, F3: 1.3 M, 1.1713 g/cm^3^, F4: 2.5 M, 1.3163 g/cm^3^) and ultracentrifuged for 3 hours at 180,000 × g (average) at 4°C (TH-641 rotor, Thermo Fisher, thinwall polypropylene tube with 13.2 ml capacity) to separate EVs and other particles based on density. After the spin, F1 (1.2ml above F2), F2 and F3 were collected, diluted with PBS, and spun for 70 min at 110,000 x g (average) at 4°C (TH-641 rotor, Thermo Fisher, thinwall polypropylene tube with 13.2 ml capacity) to collect EVs. F2 was defined previously as the EV-enriched fraction ^13^. Supernatant was discarded, and the pellet was resuspended in 100 μl PBS.

### Size Exclusion Chromatography followed by Ultracentrifugation/Ultrafiltration (SEC and UC / UF)

For SEC, the sEV-containing supernatant from the 10K step was concentrated with a 100 kilodalton (kDa) molecular weight cut-off (MWCO) protein concentrator (Thermo Fisher 88524) from 5 ml to 0.5 ml before application onto qEV Original SEC columns (IZON Science SP1-USD, Christchurch, New Zealand) that had been pre-rinsed with 15 ml PBS. 0.5 ml fractions were collected by elution with PBS. The first 3 ml (F1-6) eluate was considered the void volume, and a total of 2 ml eluate (Fractions 7-10) were collected and pooled as EV-enriched fractions. The total collection time from SEC columns was around 15 min per sample. To further purify and concentrate EVs, either ultracentrifugation or ultrafiltration was used. Ultracentrifugation was for 70 min at 110,000 x g (average) at 4°C (TH-641 rotor, Thermo Fisher, thinwall polypropylene tube with 13.2 ml capacity). Supernatant was removed, and the pellet was resuspended in 100 μl PBS. Ultrafiltration was through a 10 kDa MWCO protein concentrator (Thermo Fisher 88516), concentrating the original 2 ml to 100 μl.

### Brain homogenate preparation

For protein extraction, brain was processed to brain homogenate (BH) or brain homogenate with collagenase treatment (BHC) by grinding in cold PBS containing PI/PS with a handheld homogenizer (Kontes Pellet Pestle Motor) for 10 s. RIPA lysis buffer (Cell Signaling Technology 9806) was added, and the mixture was sonicated on ice for 2 min. Homogenate was rotated at 4°C for 2 h and spun 15 min at 14,000 x g at 4°C. Protein supernatant was transferred to tubes and stored at −80°. For RNA extraction, Trizol (Thermo Fisher 15596018) was added to frozen brain tissue and homogenized with Lysing Matrix D beads (MP Biomedicals 116913100) using a tissue homogenizer (FastPrep-24, MP Biomedicals).

### Nanoparticle Tracking Analysis

Particle concentration was measured in scatter mode (488 nm laser) with Particle Metrix ZetaView QUATT® and ZetaView® software version 8.05.10 (Particle Metrix, Germany). EV samples were diluted to a volume of 1 ml and injected into the viewing chamber by syringe. Light scattering was then recorded for 2 minutes with the following settings: Focus: autofocus; Camera sensitivity for all samples: 80.0; Shutter: 70; Scattering Intensity: 4.0; Cell temperature: 25°C. Analysis was done with parameters: Maximum particle size 1000; Minimum particle size 5; Minimum particle brightness 20.

### NanoFCM flow analysis

Particle size distribution was assessed by nanoFCM flow nano-Analyzer (NanoFCM Co.). Single photon counting avalanche photodiodes (APDs) were used for detection of side scatter (SSC) of individual particles. The instrument was calibrated for concentration and size using 200 nm polystyrene beads and a silica nanosphere cocktail (provided by NanoFCM as pre-mixed silica beads with diameters of 68, 91, 113, and 151 nm), respectively. EV preparations resuspended in 50 μl of PBS were passed by the detector and recorded for 1 minute. Using the calibration curve, the flow rate and side scattering intensity were converted into corresponding particle number and size.

### Transmission Electron Microscopy

EV preparations (10 µL) were adsorbed to glow-discharged 400 mesh ultra-thin carbon coated grids (EMS CF400-CU-UL) for two minutes, followed by 3 quick rinses in TBS and staining in 1% uranyl acetate with 0.05 Tylose. After being aspirated and dried, grids were immediately observed with a Philips CM120 instrument set at 80 kV, and images were captured with an AMT XR80 high-resolution (16-bit) 8-Megapixel camera.

### Western Blotting

EV-containing fractions were lysed in 1X RIPA lysis buffer. Protein concentrations were determined by BCA protein assay kit (Thermo Fisher). Equivalent total protein amounts from BH and EVs were separated on 4−15% stain-free pre-cast SDS-PAGE gradient gels (Bio-Rad) under non-reducing conditions and transferred onto PVDF membranes (Sigma Aldrich). After 1 hour blocking in 5% non-fat milk solution (Bio-Rad 170-6404) at room temperature, membranes were incubated with anti-CD63 (1:1000 dilution), anti-Bip (1:1000 dilution) (BD Biosciences 556019 and 610978, respectively), anti-CD81(1:1000 dilution), anti-Rab27 (1:1000 dilution) (Santa Cruz Biotechnology sc23962, sc74586), anti-TSG101 (1:500 dilution), anti-CD9 (1:500 dilution), anti-Syntenin (1:500 dilution), anti-Calnexin (1:2000 dilution), or anti-GM130 (1:1000 dilution) (the last five antibodies were Abcam ab125011, ab92726, ab133267, ab22595, and ab76154) overnight at 4°C. The membrane was washed 3 times for 8 minutes in PBST while shaking, then incubated with HRP-conjugated secondary antibody (1:10000 dilution) (Santa Cruz Biotechnology sc-2357, sc-516102) at room temperature for 1 hour. After washing again in PBST, the enzyme-linked antibody was detected by incubation with Pico chemiluminescent substrate (Thermo Fisher 34580) and recording on film (Millipore Sigma GE28-9068-38).

### RNA extraction and quality control

RNA was extracted by miRNeasy Mini Kit (Qiagen 217004) according to the manufacturer’s instructions. EV small RNA size profiles were analyzed using capillary electrophoresis by RNA 6000 Pico Kit (Agilent Technologies 5067-1513) on a Fragment Analyzer (Advanced Analytical). Total RNA and small RNA from BH were analyzed using capillary electrophoresis by RNA 6000 Nano Kit (Agilent Technologies 5067-1511) and RNA 6000 Pico Kit.

### Small RNA sequencing

Small RNA libraries were constructed from 50 ng of RNA extracted from brain homogenate or 5 µl of RNA from bdEVs using the Ion Total RNA-Seq Kit V2 (Life Technologies 4475936). Barcoding was performed with the indices from the Ion Xpress™ RNA-Seq Barcode 1-16 Kit (Life Technologies 4471250) according to the manufacturer’s protocol and as previously published^28^. The yield and size distribution of the small RNA libraries were assessed using the Agilent 2100 Bioanalyzer™ instrument with DNA 1000 chip (Agilent Technologies 5067-1504). Libraries were prepared for deep sequencing using the Ion Chef system (Life Technologies 4484177) and sequenced on the Ion Torrent S5™ using Ion™ 540 chips (Life Technologies A27765).

### Sequencing data analysis

Original BAM files were converted into FASTQ format using picard tools (SamToFastq command). Reads shorter than 15 nt were removed from the raw FASTQ using cutadapt software v1.18. The size-selected reads were aligned to human reference transcriptomes using bowtie software (1 mismatch tolerance) in a sequential manner. Specifically, reads were first mapped to rRNA, tRNA, RN7S, snRNA, snoRNA, scaRNA, VT-RNA, Y-RNA as well as the mitochondrial genome. All reads which did not map to the above RNA species were aligned to human miRNA reference (miRBase 22 release). The remaining reads were further aligned to protein-coding mRNAs and long non-coding RNA (lncRNA) references (GENCODE Release 29). The numbers of reads mapped to each RNA type were extracted using eXpress software based on a previous publication ^29^. miRNAs identified with at least 5 reads were used for further analysis. miRNA reads were normalized as reads per million miRNA reads (RPM). Differential gene expression was quantified using R/Bioconductor packages edgeR and limma as described^30^. Hierarchical clustering of miRNAs was performed with Heatmapper^31^.

### Mass spectrometry

Samples were resuspended in 1 X RIPA buffer (20mM Tris-HCl pH7.5, 150mM NaCl, 1mM Na2EDTA, 1mM EGTA, 1% NP-40, 1% SDS, 2.5mM sodium pyrophosphate, 1mM β-glycerophosphate, 1mM Na3VO4, 1ug/ml leupeptin) and protease inhibitors and incubated on ice for 5 min. The samples were then sonicated for 15 min in an ice water bath before centrifugation at 14,000g at 4°C for 10 min. Protein concentration or the supernatant was determined by Micro BCA assay (Thermo Fisher Scientific 23235). Brain homogenate (3 µg for human, 2 µg for macaque) and 10K pellet and EV samples (2 µg for human, 1 µg for macaque) were buffer exchanged to remove detergent. Protein was resuspended in 8M Urea, 50 mM Tris pH=8.3. 1 µL of TCEP (tris [2-carboxyethyl] phosphine hydrochloride, 200 mM solution in water) was then added to the samples and incubated for 4 hours at 21°C in a ThermoMixer (Eppendorf AG). 4 µL of 1M IAA (iodoacetamide in water) was then added, and samples were incubated in the dark at 21°C. 800 µL of 50 mM Tris (pH 8.3) and 1 μg trypsin were then added to samples prior to overnight incubation at 37°C. 10 μL of 10% trifluoroacetic acid (TFA) was added to each sample to acidify. Samples were cleaned using stage-tips preparations using 3 plugs of Empore polystyrenedivinylbenzene (SBD-XC) copolymer disks (Sigma Aldrich, MO, USA) for solid phase extraction.

Peptides were reconstituted in 0.1% formic acid and 2% acetonitrile and loaded onto a trap column (C18 PepMap 100 μm i.d. × 2 cm trapping column, Thermo Fisher Scientific) at 5 µL/min for 6 min using a Thermo Scientific UltiMate 3000 RSLCnano system and washed for 6 min before switching the precolumn in line with the analytical column (BEH C18, 1.7 μm, 130 Å and 75 μm ID × 25 cm, Waters). Separation of peptides was performed at 45°C, 250 nL/min using a linear ACN gradient of buffer A (water with 0.1% formic acid, 2% ACN) and buffer B (water with 0.1% formic acid, 80% ACN), starting from 2% buffer B to 13% B in 6 min and then to 33% B over 70 min followed by 50% B at 80 min. The gradient was then increased from 50% B to 95% B for 5 min and maintained at 95% B for 1 min. The column was then equilibrated for 4 min in water with 0.1% formic acid, 2% ACN. Data were collected on a Q Exactive HF (Thermo Fisher Scientific) in Data Dependent Acquisition mode using m/z 350–1500 as MS scan range at 60 000 resolution. HCD MS/MS spectra were collected for the 7 most intense ions per MS scan at 60 000 resolution with a normalized collision energy of 28% and an isolation window of 1.4 m/z. Dynamic exclusion parameters were set as follows: exclude isotope on, duration 30 s and peptide match preferred. Other instrument parameters for the Orbitrap were MS maximum injection time 30 ms with AGC target 3 × 106, MSMS for a maximum injection time of 110 ms with AGT target of 1 × 10^5^.

### Proteomics data analysis

Protein sequence data for human (last modified date: 16 May 2019) and pigtailed macaque (last modified date: 26 October 2018) were downloaded from the Uniprot database and used as the database for the search engine. Common Repository of Adventitious Proteins (CRAP) was used as the potential lab contaminant database. Protein identification was performed using the proteomics search engine Andromeda built in to Maxquant V 1.16.0. Trypsin with a maximum of two missed cleavages was used as the cleavage enzyme. Carbamidomethyl of cysteine was set as fixed modification and oxidation of methionine was set as variable modification. The Percolator results were set to reflect a maximum of 1% false discovery rate (FDR). The Label Free quantification was done with match between runs using a match window of 0.7 min. Large LFQ ratios were stabilized to reduce the sensitivity for outliers. For human datasets, data normalization was done using the Cyclicloess method. For pigtailed macaque, LFQ values were normalized using the delayed normalization described in Cox et al.^32^.

Tissue expression data were retrieved using the Database for Annotation, Visualization, and Integrated Discovery (DAVID) ^33^, while the cellular component annotations of identified proteins were enriched by Funrich^34^ and STRING^35^. Kyoto Encyclopedia of Genes and Genomes (KEGG)^36^ and Reactome^37^ was used to enrich pathway involvement of identified proteins. Statistical significance of enrichment was determined by the tools mentioned above. Only significant categories (FDR-corrected p value < 0.01) were included for analysis.

### Statistical analysis

Statistical significance of differences in total EV particle concentration, protein, and particle/protein ratio harvested from different combination of protocols were assessed by two-tailed Welch’s T test.

### Data availability

Nucleic acid sequencing data have been deposited with the Gene Expression Omnibus, accession <u>GSE150460</u>. Proteomics data files are available on request.

## Results

### Protocol comparison: bdEV separation

We followed the tissue processing and bdEV separation protocol previously published by several members of the author team (Vella et al., JEV, 2017)^13^ through the 2,000 x g centrifugation step (Figure 1). The EV-containing supernatant was filtered for stringent removal of debris, followed by 10,000 x g centrifugation. The pellet of this centrifugation step was resuspended and retained as the “10K” fraction. It should noted that this 10K fraction contains EVs of various sizes, but that purity might be low and that some large EVs were likely removed by the prior filtration. The sEV-containing 10K supernatant was then subjected to SDGU (as previously published) or SEC. Where indicated, SEC fractions were then concentrated by UC or ultrafiltration (UF) (Figure 1). Throughout this report, we will refer to the EV-containing 10K pellet as “10K”, while the final, sEV-enriched fractions that have been subjected to additional separation are referred to simply as “EVs”. The table in Figure 1 shows the source species and separation method used in all figures of the paper.

**Figure 1.**
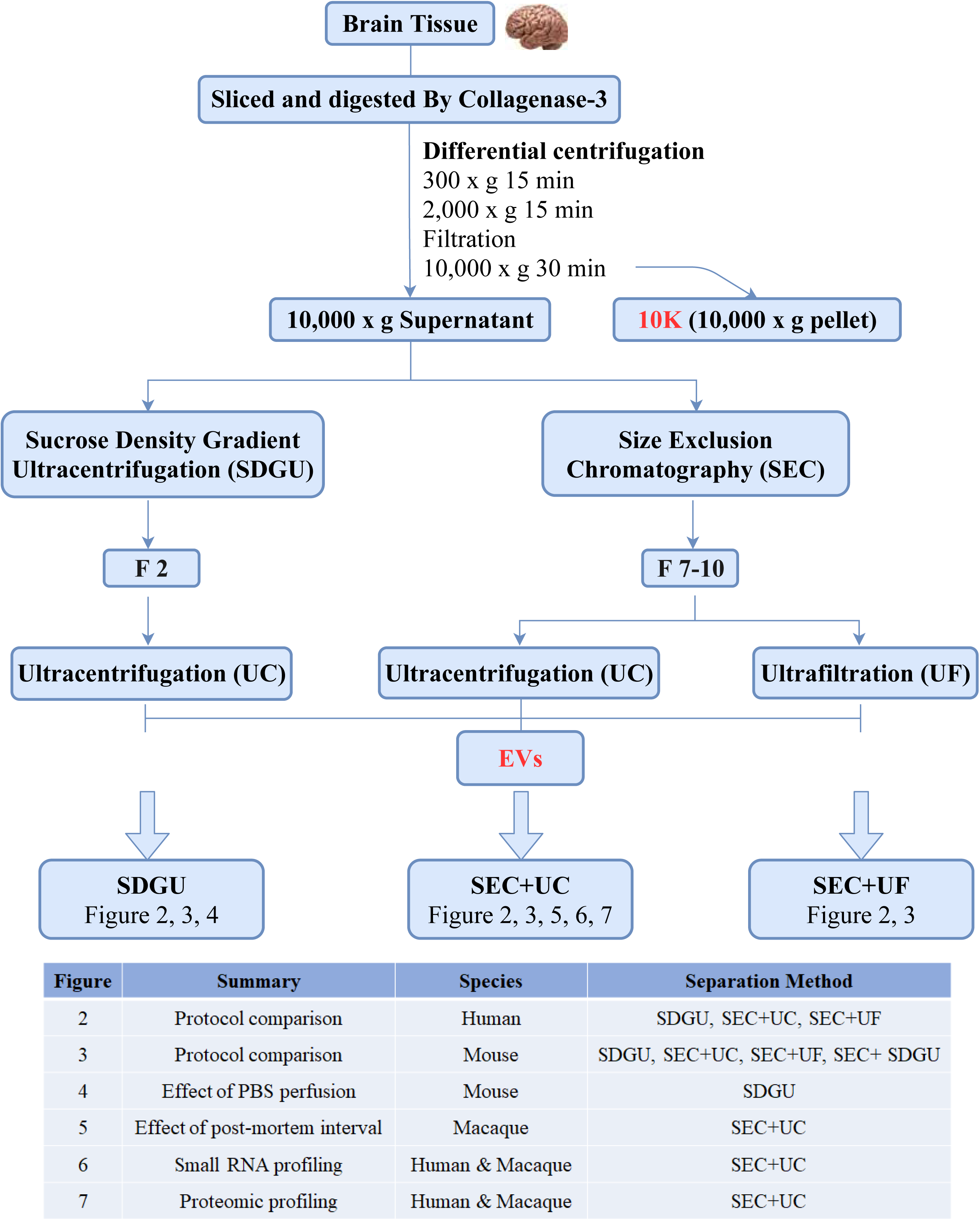
Study design and workflow for brain tissue-derived EV (bdEV) enrichment. Following digestion and centrifugation/filtration steps, 10,000 x g pellets were collected and defined as the 10K fraction. Sucrose density gradient ultracentrifugation (SDGU) or size-exclusion chromatography (SEC) were applied to 10,000 x g supernatants to enrich bdEVs from human, mouse, and macaque tissues as indicated in the table.

### Comparison of bdEV particle count, morphology, and protein markers in different fractions of SDGU and SEC: human and mouse tissue

Particle yield per 100 mg tissue input (human or mouse) was determined by nanoparticle tracking analysis (NTA). Particle yield was highest for F2 from SDGU and F7-10 from SEC+UC compared with other fractions. Particle concentration was below the reliable range of measurement in F3-6 and F11-14 from the SEC+UC method. Particle yield was similar for human and mouse brains (Figure S1A-B). Transmission electron microscopy (TEM) revealed cup-shaped oval and round particles in SDGU fractions (Figure S1C) and F7-10 from SEC+UC (Figure S1D) that were consistent with EV morphology. Presence of EV-enriched membrane (CD9, CD63, CD81) and cytosolic (TSG101, syntenin) markers, and expected EV-depleted cellular markers (GM130, calnexin, BiP) was examined by Western blot for EV fractions as well as brain homogenate (BH), including BH treated with collagenase (BHC). For human brain-derived EVs, abundant CD63 and CD81 levels were observed in F2 and F3 from SDGU and F7-10 from SEC+UC, while F11-14 (SEC) were positive for CD63 but not CD81. Compared with more purified, smaller EVs, the 10K fraction had lower but still detectable CD63 and CD81. bdEVs from human samples generally did not have detectable calnexin or GM130, in contrast with the source BH and BHC (Figure S1E-F). However, mouse bdEVs retained some amount of cellular markers regardless of separation technique (Figure S1G-H). Additionally, some apparent differences between EV markers in the different fractions were also observed between human and mouse.

### Characterization of human bdEVs obtained by three combinations of methods

Based on the results above, we focused on F2 of SDGU and F7-10 of SEC and examined the output of SDGU compared with SEC+UC and SEC+UF (see sample information, Table 1). In these experiments, NTA suggested that the SEC methods yielded a slightly higher number of particles as compared with SDGU (Figure 2A). There was no significant protein concentration difference of EVs obtained by the three methods (Figure 2B), but particle:protein ratio, often used as a surrogate of EV preparation purity, was highest for the SEC + UC methods compared with SDGU (Figure 2C). TEM revealed particles consistent with EV morphology from all methods (Figure 2D). In terms of putatively EV-enriched and -depleted markers, all methods were consistent with EV marker enrichment and cellular marker depletion (Figure 2E). A small amount of calnexin was found in an SEC+UF lane, though, possibly indicating retention of some fragments by UF that are cleared by UC. We concluded from these results that, following SEC, UC may have a slight advantage in increasing purity compared with UF. The 10K fraction had lower particle concentration but higher protein concentration compared with more extensively purified EVs obtained from SEC, resulting in low particle:protein ratios (Figure 2 A-C). TEM showed presence of EVs in the 10K fraction (Figure 2D) while WB showed less CD63 and CD81 in 10K than in more purified EVs (Figure 2E).

**Figure 2.**
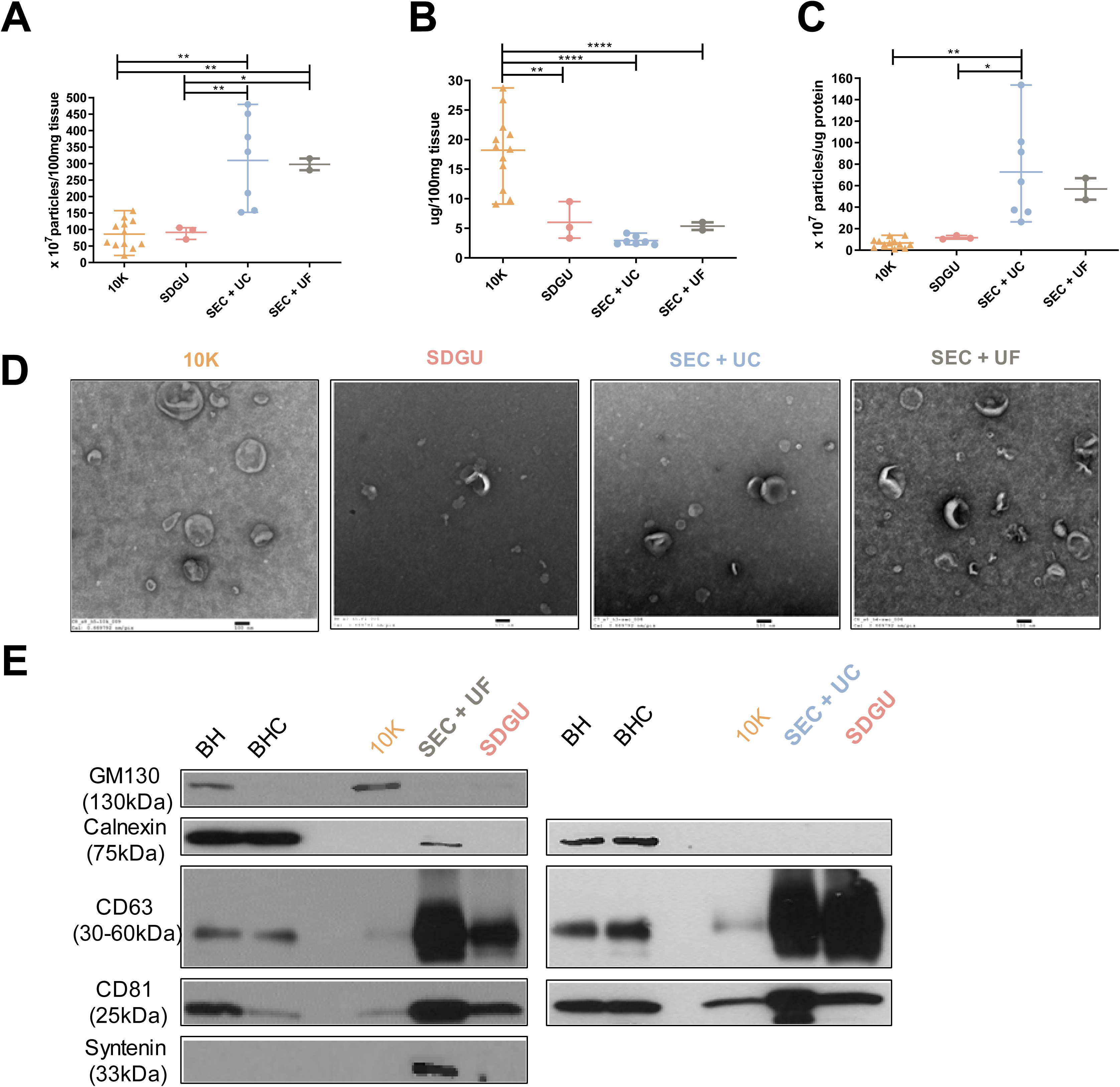
Characterization of human bdEVs obtained by three combinations of methods. (A) Particle concentration of human bdEVs separated by three combinations of methods was measured by NTA (Particle Metrix). Particle concentration for each group was normalized by tissue mass (per 100 mg). (B) Protein concentration of human bdEVs separated by different methods and measured by BCA protein assay (per 100 mg tissue). (C) Ratio of particles to protein (particles/µg). (A)-(C): Data are presented as the mean with range. *p ≤ 0.05, **p ≤ 0.01, ***p ≤ 0.001, ****p ≤ 0.0001 by two-tailed Welch’s t-test. (D) bdEVs were visualized by negative staining transmission electron microscopy (scale bar = 100 nm). TEM is representative of five images taken of each fraction from three independent human tissue samples. (E) Western blot analysis of GM130, calnexin, CD81, CD63, and syntenin associated with BH and EV fractions. WB are representative of three independent human tissue EV separations from the SDGU and SEC+UC methods, and one independent human tissue EV separation from the SEC+UF method.

### Comparison of methods for mouse bdEV separation

We next applied the same methods to perfused mouse brain (see sample information, Table 2). Since our preliminary results indicated more contamination of mouse bdEVs with cellular markers (Figure S1 G-H), we also added a fourth method, in which both SEC and SDGU were used. Yield of particles was highest for the SEC+UF combination compared with SDGU and SEC+UC (Figure 3A), while protein yield was higher in SEC+UF and SDGU compared with SEC+UC (Figure 3B). Purity, as estimated by particle:protein ratio, was highest for SEC+UF, then SEC+SDGU and SEC + UC, while SDGU was the lowest (Figure 3C). However, particles obtained by SEC+UF included more non-vesicular material, suggesting again that concentration by UC also served a purification function relative to UF (Figure 3D). Cellular markers GM130, Calnexin, and Bip were found despite method (Figure S1G, H, Figure S2). Interestingly, in contrast with the human brain results, the particle and protein concentrations of 10K fractions were both higher than those of more purified EVs, contributing to a moderate particle/protein ratio compared with EVs (Figure 3A-C). As expected, 10K also contained EVs by TEM (Figure 3D) and included a large number of cellular proteins (Figure S1G,H, Figure S2).

**Figure 3.**
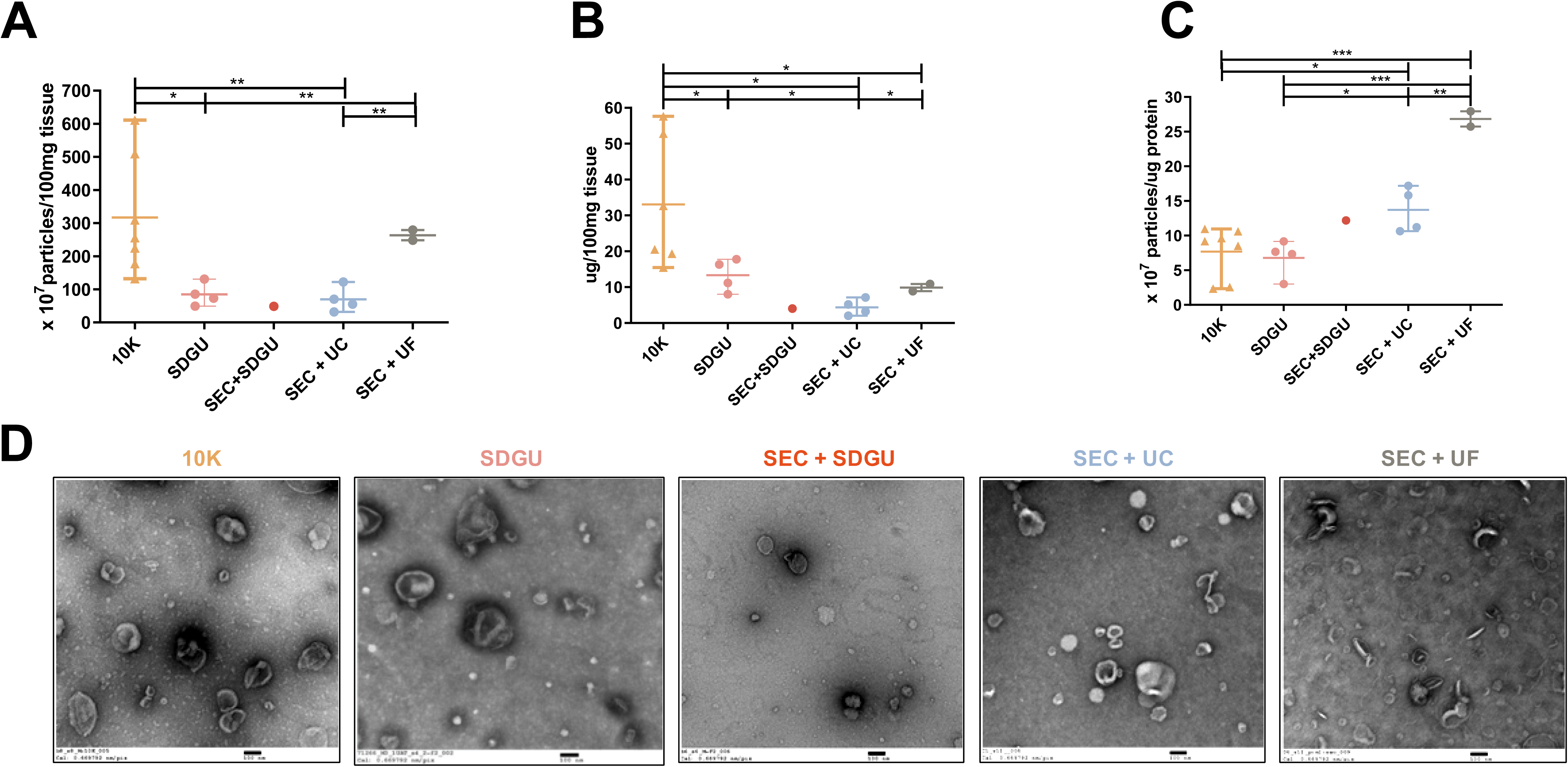
Characterization of mouse bdEVs. (A) Particle concentrations of mouse bdEVs separated by different methods were measured by NTA (Particle Metrix, normalized per 100 mg tissue). (B) Protein concentration of mouse bdEVs separated by different methods and measured by BCA protein assay kit (per 100 mg tissue). (C) Ratio of particles to protein (particles/µg). (A)-(C): Data are presented as the mean with range. *p ≤ 0.05, **p ≤ 0.01, ***p ≤ 0.001, ****p ≤ 0.0001 by two-tailed Welch’s t-test. (D) bdEVs were visualized by negative staining transmission electron microscopy (scale bar = 100 nm). TEM is representative of five images taken of each fraction from three independent tissue samples.

### Does PBS perfusion at necropsy affect bdEV separation?

Considering the large amount of cellular markers detected in bdEV preparations from mouse, we reasoned that tissue preparation might be altered to reduce this influence. For example, in animal models, it is often possible to perfuse with buffer (such as PBS) at necropsy, flushing blood from the tissues. We thus separated bdEVs from tissue of animals perfused or not with PBS (the sample information sees Table 2, mouse 1-4 and 12-15) . Although no remarkable differences were observed for particle yield (Figure 4A), perfusion was associated with apparent depletion of the Golgi marker GM130 (Figure 4B). However, presence of calnexin and BiP was similar, and apparent changes in some EV-associated proteins were observed (Figure 4B).

**Figure 4.**
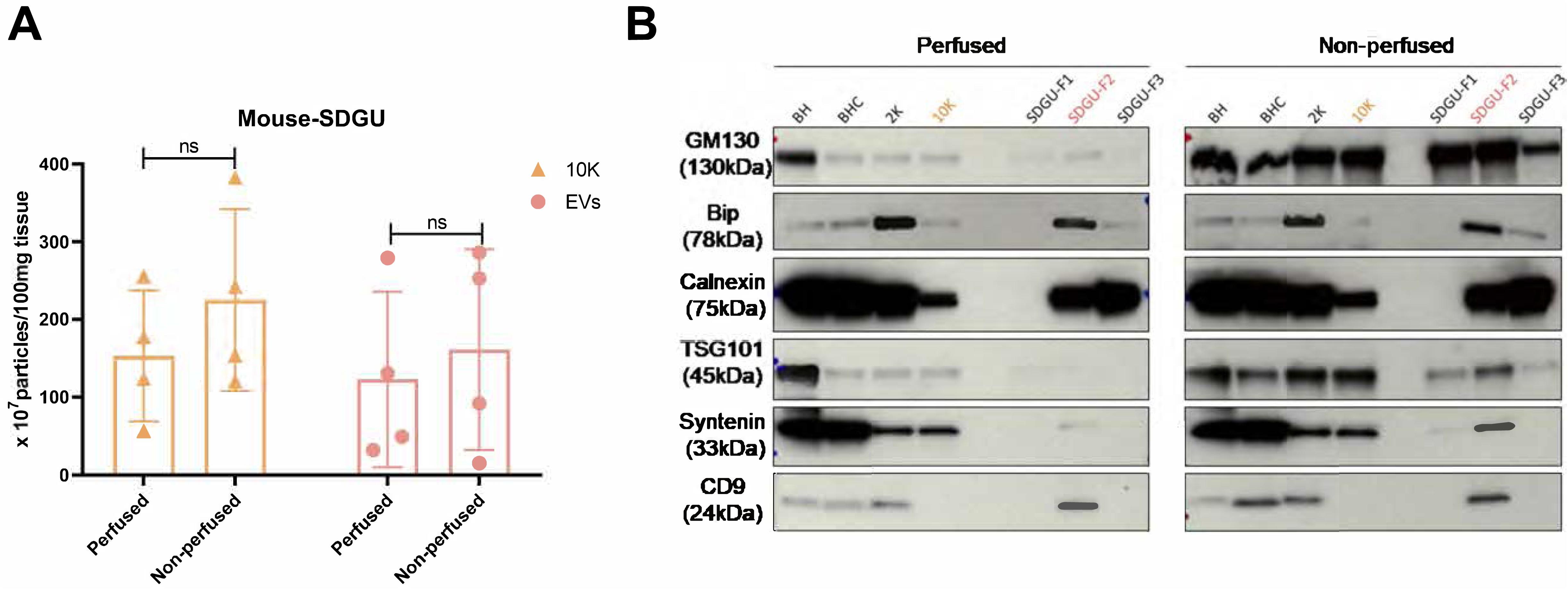
Effect of PBS perfusion on mouse bdEV separation. (A) Particle concentration of mouse bdEVs from PBS-perfused and non-perfused brains as separated by SDGU. Data are presented as the mean with range (n = 4). ns: No significant difference was detected between perfused and non-perfused samples by two-tailed Welch’s t-test. (B) Western blot analysis of GM130, Bip, calnexin, TSG101, syntenin, and CD9 associated with BH and EV fractions from perfused and non-perfused mouse brains. Blots are representative of three independent mouse bdEV separations using SDGU.

### What is the effect of post-mortem interval on bdEV separation?

Whereas brain tissue from animal models such as mouse can be processed (or snap-frozen and stored) immediately after necropsy, human brains are acquired, cut, stored, and/or otherwise processed after varying times and temperatures. To assess the influence of one part of the length of time between death and processing (post-mortem interval, PMI), specifically the time after sectioning and before final bdEV extraction, we obtained occipital lobe of macaques that were sacrificed in the course of other studies (see sample information, Table 3). Lobes from the same subject were divided into three parts and placed at room temperature for two, six, or 24 hours (2, 6, 24H). bdEVs were then obtained using the method ending with SEC+UC as outlined above. bdEVs from tissue incubated for 24H had a higher particle yield compared with 2H and 6H (Figure 5A). By TEM, bdEV morphology was the same for macaque as for human and mouse EVs (Figure 5B). In contrast, particle concentration of the 10K fraction did not appear to be affected by the investigated PMI time points (Figure 5A).

**Figure 5.**
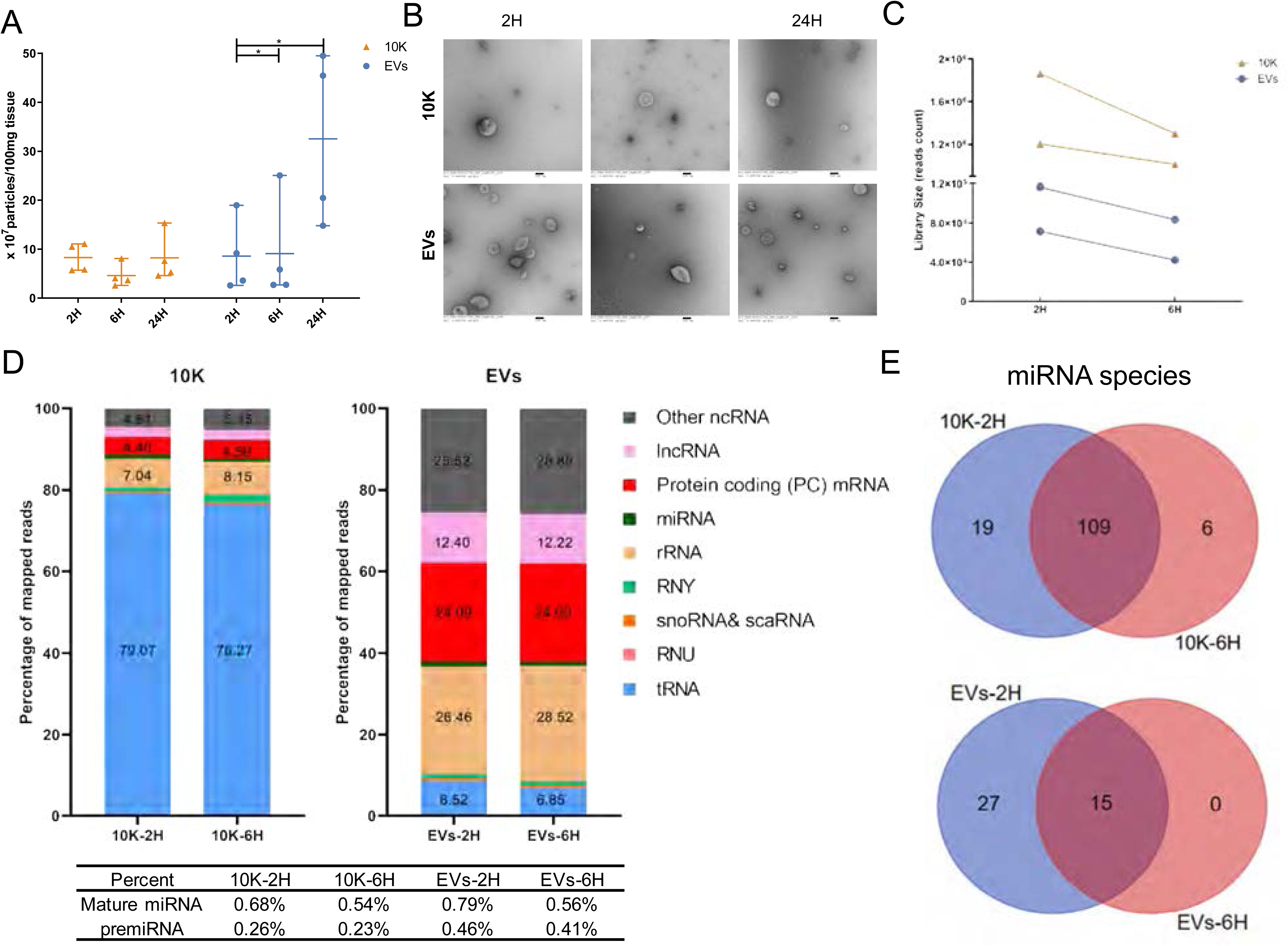
Effect of postmortem interval on macaque bdEV separation. (A) Particle concentration of bdEV preparations by SEC+UC from 2H, 6H and 24H postmortem interval (PMI) macaque brains was measured by NTA (Particle Metrix, normalized per 100 mg tissue). Data are presented as the mean with range (n=4). *p ≤ 0.05, **p ≤ 0.01, ***p ≤ 0.001, ****p ≤ 0.0001 by two-tailed Welch’s t-test.(B) BdEVs from 2H, 6H and 24H PMI macaque brains were visualized by negative staining transmission electron microscopy (scale bar = 100 nm). TEM is representative of five images taken of each fraction from two independent tissue samples. (C) Small RNA sequencing; total reads from 10K and EVs from 2H and 6H PMI (n=2). (D) Average percent (n=2) of mapped reads for the nine most abundant RNA classes in 10K and EVs at 2H and 6H PMI. (E) Venn diagram of miRNAs (mean raw reads > 5) from two independent 10K and EV preparations at 2H and 6H PMI.

### Effect of PMI on bdEV small RNA and protein content

Ligation-dependent small RNA sequencing was used to assess the effects of PMI on bdEV small RNA content from tissues of two macaques (Table 3, macaques 3 and 4). Only 2H and 6H libraries were sequenced because quality control electropherograms of sequencing libraries prepared from 24H PMI samples suggested some level of degradation (Figure S3). The mapped read counts were also slightly lower at 6H compared with 2H for both 10K fractions and more extensively separated EVs from two macaques (Figure 5C). Strikingly different proportions of RNA biotypes were detected for 10K and EVs, but with minimal differences from 2H to 6H (Figure 5D). Prominently, more purified EVs fractions contained a smaller percentage of tRNA-related sequences than 10K fractions. In terms of miRNA diversity, “abundant” miRNAs were defined inclusively as those with greater than 5 counts. In 10K and EVs, most miRNAs were found at both 2H and 6H, whereas more miRNAs were exclusively detected at 2H (Figure 5E). These findings suggest that miRNA diversity may decrease somewhat as PMI increases.

Because of limited input of macaque protein for proteomics (1 µg), only limited numbers of proteins were identified in either the 10K or more purified, sEV-enriched EV fractions from the qualitative proteomics analysis (Figure S4 A). The number of identified proteins increased for 10K and EVs with greater PMI, while it appeared to decrease in BH (Figure S4 A). Almost all 10K and EV proteins identified at 2H and 6H were also identified at 24H. Many proteins were found only at 24H. 75% of proteins were found at all time points in BH. Moreover, in both BH and the two EV-containing fractions, the overlap of proteins at 6H and 24H was greater than that for 2H and 24H, indicating a time-dependent shift in protein detection. Based on pathway involvement, proteins identified in EVs were especially enriched for known EV (“extracellular exosome”), cytoplasmic, lysosomal, and membrane-associated proteins (Figure S4 B). A stronger enrichment of cytoplasmic components at 24H might indicate more cell disruption in these samples (Figure S4 B). 10K shared many components with more purified or smaller EVs, but were also enriched for terms such as desmosome and centrosome that did not appear in the EV fractions. Importantly, though, the meager protein coverage from these samples limits the conclusions we can draw from the effect of PMI on EV protein content. We also validated the EV-enriched and - depleted markers. Both EV (CD63, CD81, TSG101, syntenin) and cellular (calnexin) markers were higher at 24H in EVs (Figure S4C), while CD63 and calnexin were lower at 24H in 10K.

### Small RNA profiling of brain-derived 10K fractions and EVs

We next examined small RNA and protein profiles of brain 10K fractions and EVs separated by the SEC+UC method from macaque and human. These included two macaque brains (Table 3, subjects 3 and 4, 2H samples) and seven human brains without recorded neurologic or cognitive abnormalities (Table 1, subjects 4-10). We examined the particle size of 10K fractions and EVs obtained by the SEC+UC method before profiling. 10K and EVs had an overlapping size distribution. More particles ranging from 40-70 nm (diameter) were detected in the sEV-enriched, more purified EV fraction, while more particles ranging from 70-145 nm were detected in the 10K fraction (Figure S5). Total RNA extraction and quality control indicated a slightly higher RNA yield from 10K compared with EVs (Figure 6A). Small RNA sequencing revealed a difference in small RNA biotype distribution between BH, 10K, and EVs (Figure 5D, Figure 6B). miRNAs, snoRNAs, and scaRNAs were more enriched in BH, tRNA fragments were more enriched in 10K, and rRNA and mRNA fragments were more enriched in EVs. Consistent with results for macaque bDEVs (as shown previously in Figure 5D), tRNA and miRNA sequences were enriched in 10K over EVs (Figure 6B). The pattern of small RNA expression was thus different not only between BH and EV-containing fractions overall, but also between 10K and EVs (Figure 6C).

**Figure 6.**
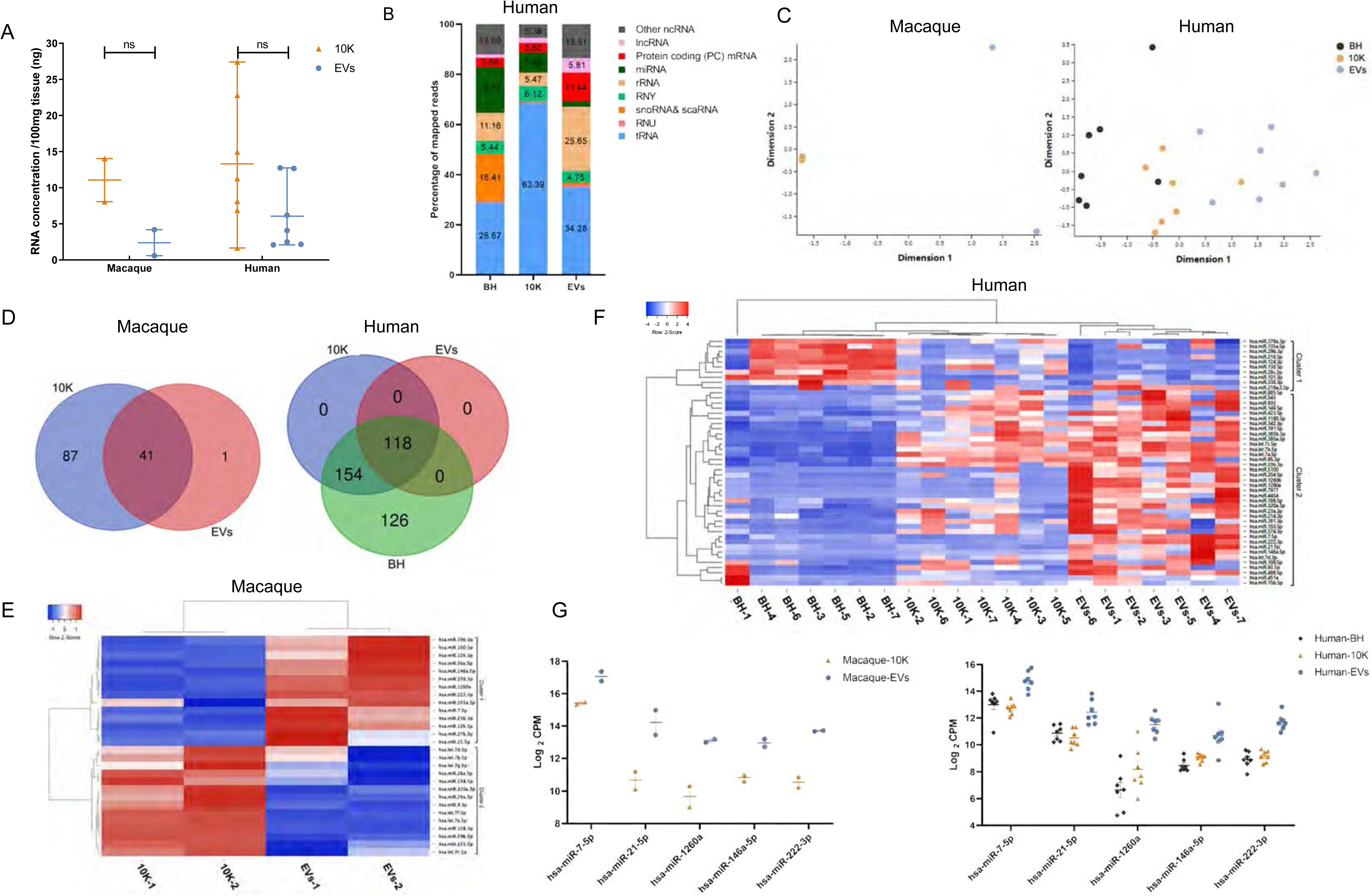
Small RNA profiles of bdEVs from two macaque and seven human brain samples. (A) RNA concentration of 10K and EV fractions prepared by SEC+UC from macaque and human brain. Data are presented as the mean with range. ns: No significant difference was detected between perfused and non-perfused samples by two-tailed Welch’s t-test. (B) Average percent of mapped reads of the nine most abundant RNA classes in BH, 10K, and EVs (human, n=7). (C) Multidimensional scaling analysis based on quantitative small RNA profiles of BH, 10K, and EVs (n=2 for macaque, n=7 for human). (D) Venn diagram of miRNAs (mean raw reads greater than 5) in 10K, EVs, and BH from macaque and human brain (n=2 for macaque, n=7 for human). (E) Unsupervised hierarchical clustering of 28 differentially abundant miRNAs of 10K and EVs (macaque, n=2). Cluster 1: enriched in EVs; Cluster 2: enriched in 10K. (F) Unsupervised hierarchical clustering of 48 differentially abundant miRNAs (BH vs EVs, mean fold change > 2; human, n=7). Cluster 1: enriched in BH; Cluster 2: enriched in EVs. (G) Abundance comparison of five miRNAs enriched in EVs vs 10K and/or BH for both primate species.

For macaque samples, numerous miRNAs were found in 10K but not in EVs, while only one miRNA appeared to be detected uniquely in EVs (Figure 6D). For human samples, 118 miRNAs were detected in common between the two EV-containing fractions and BH. All 154 miRNAs found in 10K but not EVs were also detected in brain tissue, while additional miRNAs were found in tissue but not vesicles (Figure 6D). Normalizing by CPM, among 41 macaque miRNAs found in both 10K and EVs, 28 with putative differential abundance were used for unsupervised clustering (Figure 6E, where clusters 1 and 2 are enriched in EVs and 10K, respectively). Similarly, unsupervised clustering was done with 48 human miRNAs with fold change >2 between BH and EV (Figure 6F, where clusters 1 and 2 are enriched in BH and EVs, respectively). Interestingly, but perhaps not unexpectedly, 10K have a miRNA profile different from but intermediate between BH and EVs. We were also able to identify a small minority of miRNAs that appeared to be enriched in EVs, with consistency across the two investigated species (Figure 6G).

### Proteomic profiling of brain-derived 10K and EVs

The same samples were then examined for protein concentration and profile. As expected, for both human and macaque materials, protein concentration decreased from BH to 10K and EVs (Figure 7A). For macaque samples, numerous proteins were identified in tissue only, with just a handful identified in 10K and EVs (Figure S4 A). The lower proteome coverage is likely due to the lower protein input for macaque EVs. Better coverage was obtained with human samples, for which more protein was available. Proteins identified in samples from at least five donors were included in analysis. 214 proteins were found in both the 10K and EVs. Around 215 proteins were detected only in EVs, 65 only in 10K. 99% of the detected proteome matched entries in existing EV databases Vesiclepedia^38^, EVpedia^38^, and Exocarta^39^. 71 EV proteins and 65 10K proteins were found among the top 100 most commonly reported EV proteins in these databases (Figure 7C, Table 4).

**Figure 7.**
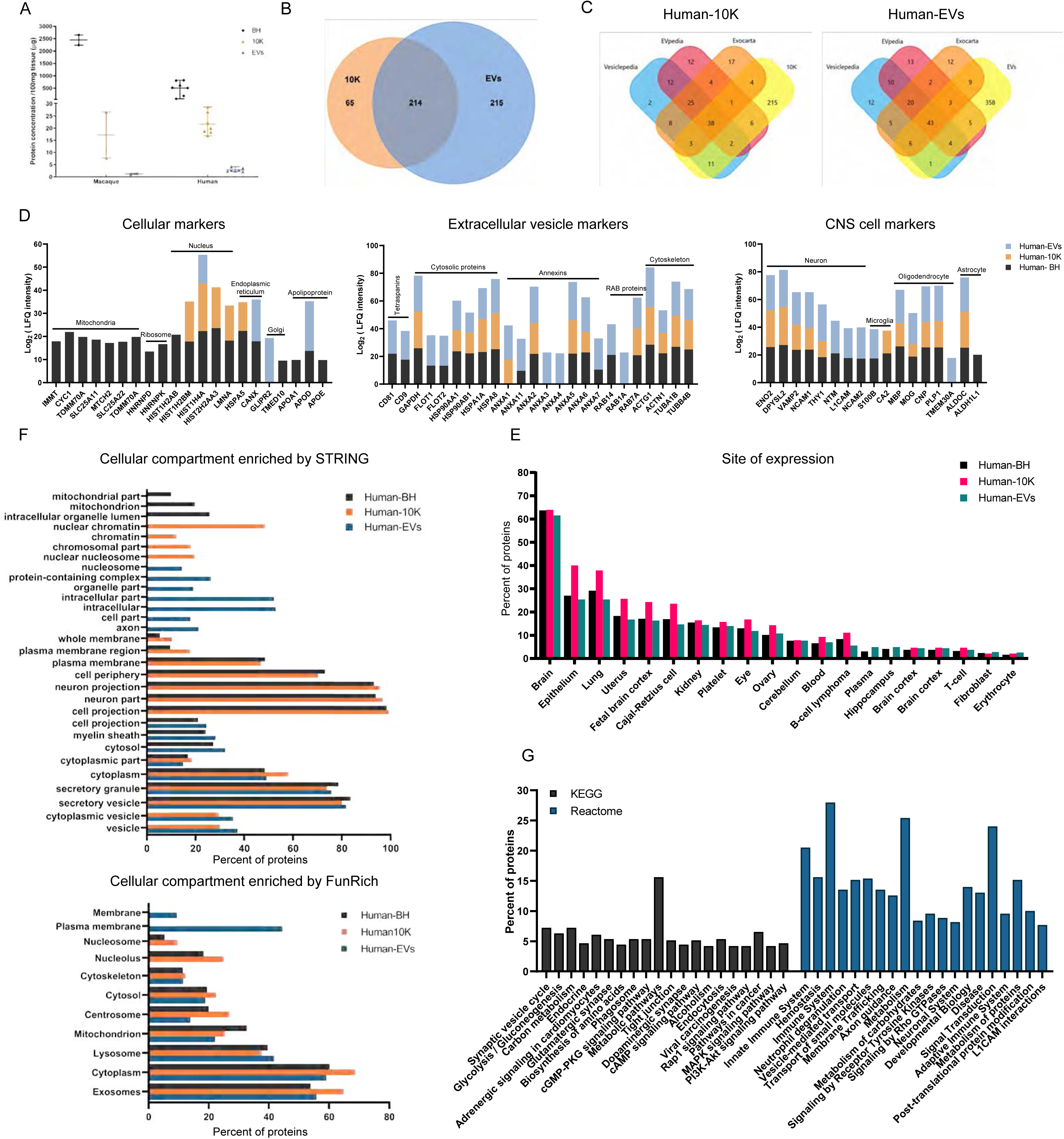
Protein profiles of EVs from two macaque and seven human brain tissues. (A) Protein concentration of BH, 10K, and EV fractions from macaque and human brain (BCA). *p ≤ 0.05, **p ≤ 0.01, ***p ≤ 0.001, ****p ≤ 0.0001 by two-tailed Welch’s t-test. (B) Venn diagram of identified proteins in 10K and EVs of human (proteins identified in 5 from n=7 individuals). (C) Ven diagram of human 10K and EV proteins matched to the top 100 proteins in EV databases Vesiclepedia, EVpedia, and Exocarta. (D) Expression levels of intracellular, extracellular vesicle (EV) and central nervous system (CNS) proteins in human brain EV, 10K, and BH preparations. (n=7) (E) Tissue derivation of human BH, 10K, and EV proteins (based on DAVID knowledgebase; the top 20 terms ranked are by detected protein number in EVs, with FDR-corrected p-value < 0.01 are shown). (F) Cellular compartments of human BH, 10K, and EV proteins (STRING and FunRich; the top GO terms are ranked by detected protein number in EVs and BH, and those with FDR-corrected p-value < 0.01 are shown). (G) Pathway involvement of human bd-sEV proteins according to the Kyoto Encyclopedia of Genes and Genomes (KEGG) and Reactome. The top 20 pathways in EVs are ranked by FDR-corrected p-value.

**Table 4.**
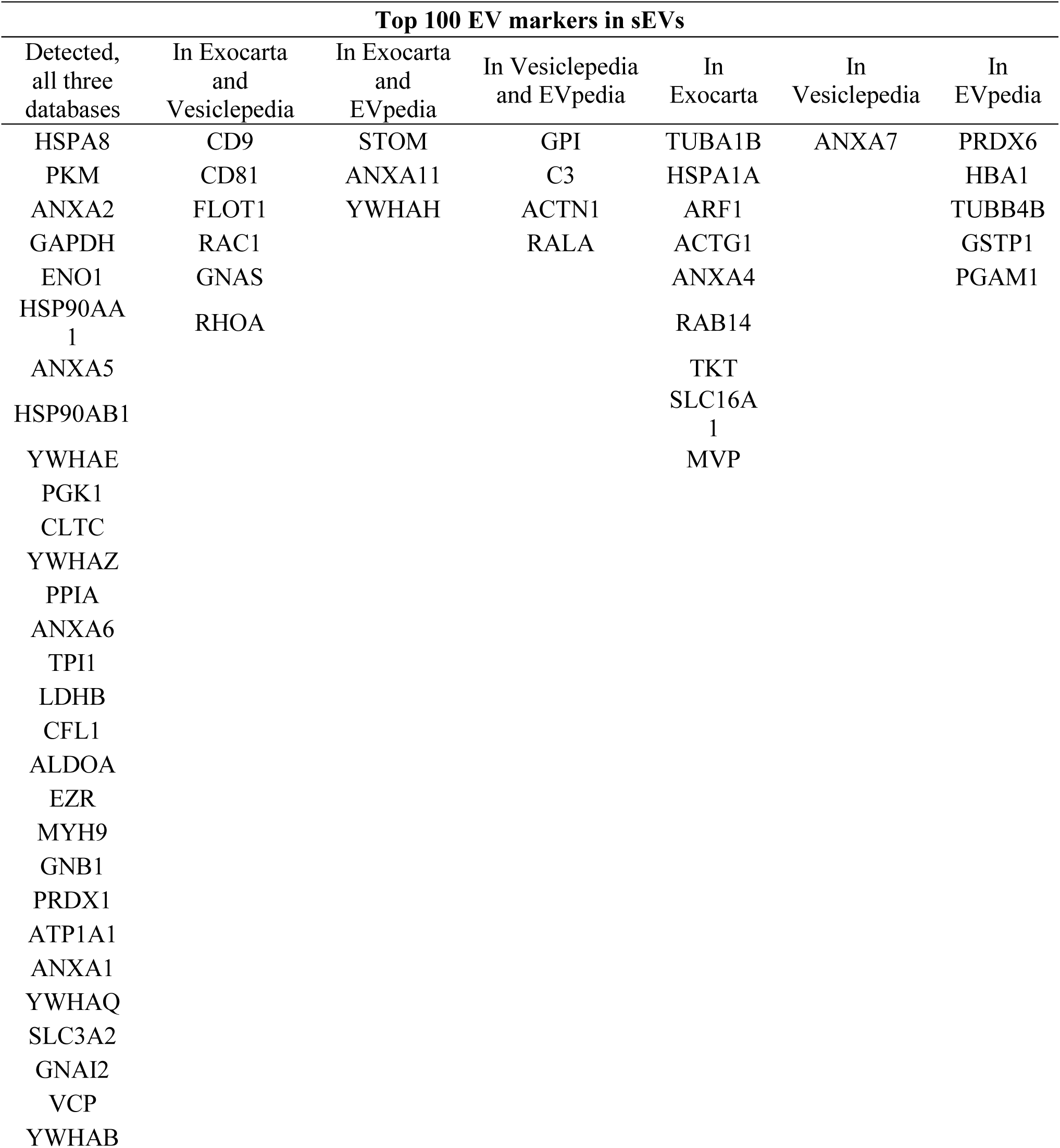
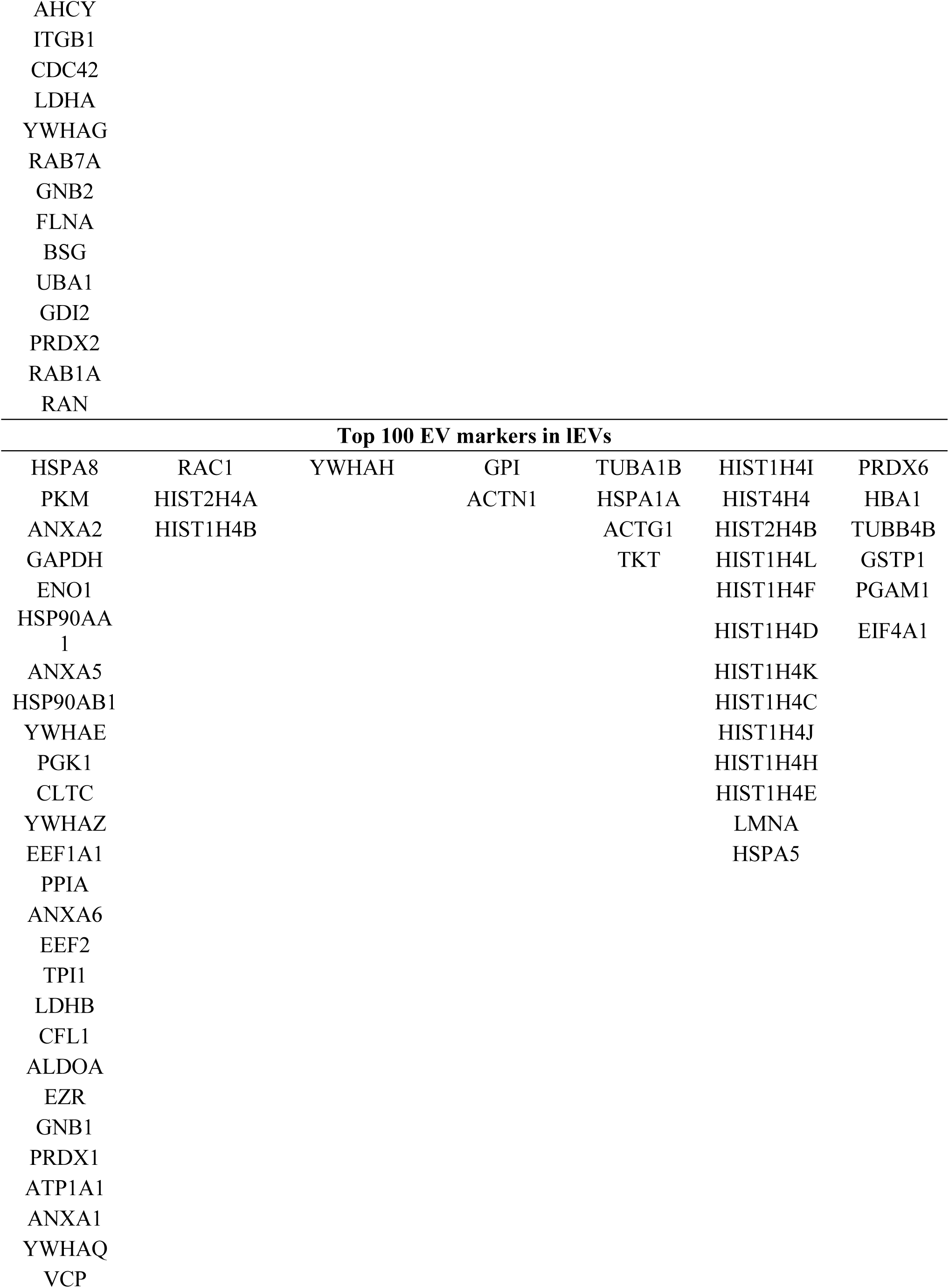
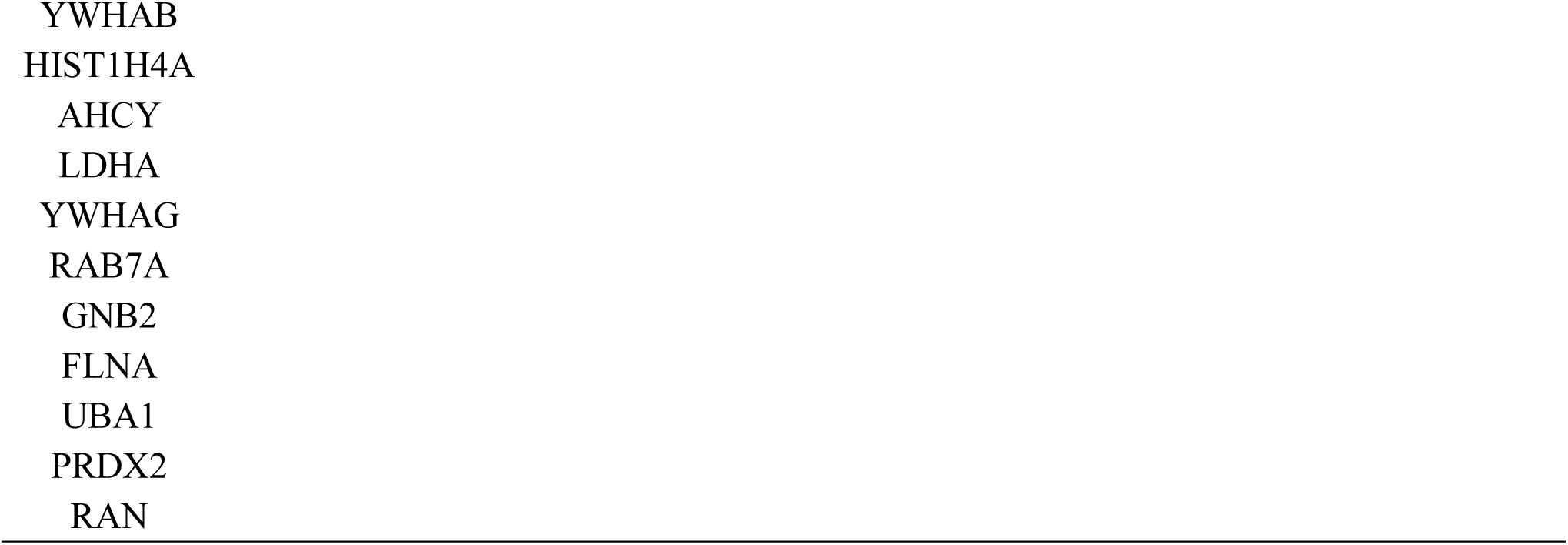
Protein list of human brain sEVs and lEVs matched to top 100 markers in Exocarta, Vesiclepedia, and EVpedia

Based on published literature and MISEV2018 suggestions, we examined known markers of cells (focusing on presumably or reportedly “EV-depleted” proteins), EVs, and the central nervous system. For the most part, presumed cellular proteins were enriched in BH, including those associated with mitochondria, ribosomes, nucleus, endoplasmic reticulum (ER), and Golgi (Figure 7D and Figure S6, left panel). Only small amounts of these proteins were found in bdEV preparations; of these, nuclear proteins were found mostly in 10K preparations, while apolipoprotein D associated with EVs (Figure 7D and Figure S6). Presumed EV proteins were found in all types of samples (Figure 7D, middle panel) but were almost all enriched in EV preparations (Figure S6, right panel). A clear enrichment of certain EV markers was observed for EVs, including tetraspanins (CD81 and CD9), cytosolic proteins (FLOT1, FLOT2), annexins (ANXA11, ANXA3, ANXA4), RABs (RAB14 and RAB1A) and cytoskeleton proteins (ACTN1). Importantly, bdEVs also carried markers of central nervous system cell types: neurons, microglia, oligodendrocytes, and astrocytes (Figure 7D, right panel). Some of these associations (e.g., for TMEM30A), may suggest selective protein packaging into EVs that could be exploited for selective enrichment of specific bdEV populations from tissue or biofluids. Presence of brain proteins was corroborated by an analysis using the DAVID database, revealing a high enrichment of brain-derived proteins in EVs (n=264), 10K (n=179) and BH (n=358) (Figure 7E). Of note, a small number of proteins enriched for terms such as platelet, blood, plasma, T-cell, fibroblast, and erythrocyte may indicate blood cell debris, infiltrating immune cells, or simply non-specificity of some proteins.

GO ontology analyses by STRING and FunRich were used to determine the cellular compartment of proteins recovered from BH and bdEV preparations (Figure 7F). Again, a large portion of proteins associated with both 10K and more purified EVs were enriched for EV-related terms like exosomes, vesicle, and cytoplasmic vesicle. The terms membrane, plasma membrane, and whole membrane, instructively, were enriched only for EVs, while nucleosome, nucleolus, protein containing complex, and intracellular part were enriched only for 10K and BH. Cytoplasm, lysosome, and cytoskeleton were common terms enriched for all groups, while some were exclusively enriched in one group. KEGG and Reactome analyses further confirmed processes related to the CNS, including synaptic vesicle cycle, axon guidance, neuronal system, and L1CAM interactions, but also identified processes related to metabolism, endocytosis, the immune system, hemostasis, vesicle-mediated transport, membrane trafficking, and signal transduction (Figure 7G).

## Discussion

Adapting the protocol previously published by Vella, et al.^13^, we evaluated application of SEC to bdEV enrichment from brain tissue. The modified method including SEC achieves a bdEV separation efficiency that may achieve acceptable results compared with SDGU. Furthermore, the modified method decreases operation time after the 10,000 x g centrifugation step from three hours (SDGU spin time) to 15 minutes (SEC on-column collection time), and the SEC step can be almost fully automated, potentially reducing operator variability. We do not suggest that SEC replaces SDGU, which may still be required when the highest levels of purity are needed. Rather, SEC may be an acceptable alternative to process larger numbers of samples or to achieve automation.

Both protocols worked well for human tissues, as assessed by the depletion of “deep cellular” markers. However, for mouse and macaque brain in particular, a substantial cellular component was still detected consistently. Since tissue preparation variables might contribute to cellular contamination, we tested for the first time the effects of tissue perfusion and postmortem interval. While perfusion had a noticeable effect in depleting cellular contaminants like GM130, cellular markers were still observed even in bdEVs from PBS-perfused mouse brain. Regarding PMI before final bdEV processing, 24 hours of tissue storage resulted in higher particle recovery, but also with a significantly higher number of intracellular proteins in macaque bdEVs. PMI did not substantially change the RNA biotypes of macaque bdEVs, but some small RNA degradation and lower miRNA diversity was associated with long PMI. On the contrary, we did not observe significant protein and RNA difference among bdEVs from human tissues with different PMI (death to acquisition), possibly indicating tissue fragility differences among species. However, we also used cortical tissues (mostly gray matter) for human, while a more varied mixture of gray matter and white matter is typically present for the much smaller structures that are available for macaque and mouse bdEV preparation. This factor could partly confound species comparisons. We conclude that perfusion is advised if possible (with animal models), and that PMI, storage temperature, and other details about tissue preparation and processing should be recorded and reported when describing bdEV separation. We note that it was recently reported that flotation density gradient separation showed lower detectable intracellular proteins compared with velocity gradients ^22^; however, flotation gradients require even more time than velocity SDGU. Since overly aggressive tissue digestion may contribute to release of cellular materials^40^, and since tissue structure may vary by species, types and concentrations of enzymes and different storage times and temperatures should be explored to find the minimal adequate digestion condition for tissues of different organisms as well as different tissue regions. We would like to point out that our study of PMI is limited. Further studies are needed to understand factors such as: temperature and time of brain in the skull after death and before removal; brain storage parameters before sectioning; and additional storage conditions before bdEV extraction.

Since the study of bdEV small RNAs is still in an early phase^13, 15^, we investigated small RNA composition of brain tissue and bdEVs. The results, with slightly different composition of 10K pellets and more separated/smaller EVs, suggested that not all small RNAs are uniformly loaded from cells into EVs. Consistent with other reports ^13, 41–45^, we also found that fragments of rRNA and tRNA, were more abundant than miRNAs in EVs, even without employing ligation-dependent sequencing library preparation^46^. Although some publications have reported a higher miRNA proportion in EVs ^47, 48^ than in cells, our results suggested that mapped miRNA reads and diversity gradually decreased from brain homogenate to 10K and then EVs. At least as assessed by our library preparation and analysis methods, miRNAs account for less than 1% of small RNA in both human and macaque bdEVs. Nevertheless, several miRNAs were highly enriched in EVs versus 10K and/or tissue. These included miRNAs (miR-423, miR-320a, miR-186, miR-146a, miR-1180, let-7b, and let-7d) that were also reported by Vella, et al., who collected data following SDGU bdEV separation ^13^. Additionally, miRNAs such as miR-7, miR-21, miR-1260, miR-146a, and miR-222, were enriched in EVs over 10K in both macaque and human datasets. Collectively, these data show that 10K and EVs, despite some likely overlap, may harbor different proportions of small RNA including miRNA, which may further differ from the ratios in parent cells. Whether this observation is explained by active packaging/exclusion or passive factors (i.e. average distance of specific RNAs from sites of EV biogenesis) remains unresolved, with evidence for and against active sorting ^41, 49^ ^42, 48, 50, 51^.

Since numerous publications have reported tissue-derived EVs without thorough protein characterization of EV-depleted cellular markers and specific EV subtype markers^12, 18, 20, 21, 52–54^, we compared the BH and bdEV proteomes with established EV-related databases. Substantial concordance between our data and the databases was clear (Tabe 4). However, reduced presence of most cellular proteins in bdEVs compared with BH indicated relatively minimal cell disruption during EV separation. In 10K, however, EV markers were found alongside higher levels of histones and other proteins that may have derived from broken nuclei or formation of amphisomes^43, 55^. Additionally, sensitive proteomics methods indicated several cellular or co-isolate markers, including calnexin, GLIPR2, and ApoD, in EVs, though we did not find Calnexin or GM130 expression by WB. These proteins that are associated with intracellular compartments beyond plasma membrane (PM)/ endosomes^56^ may be contaminants, but could also be *bona fide* cargo of bdEVs^57–61^: calnexin is carried by syncytiotrophoblast EVs with immunoregulatory function in preeclampsia^58^, while ApoD transported by astroglial cells can exert neuroprotective effects^60^. Numerous cytosolic, annexin, Rab, and cytoskeleton proteins were also found. We conclude from the endosomal and PM markers that EVs prepared by our method are a mixture of different subtypes of EVs, including true exosomes and ectosomes/microvesicles^62^, accompanied by lower amounts of proteins from other cellular compartments. In this study, the 10K pellet can be considered an intermediate fraction between BH and more highly purified and generally smaller EVs. This fraction indeed displayed more intracellular markers and fewer EV markers compared with EVs separated from the supernatant of the 10k step. Our findings are consistent with a recent proteomics study of melanoma tissue-derived EVs^16^, which showed that larger tissue EVs (in this case, from a 16.5K pellet) were more associated with intracellular markers compared with small, low-density EVs separated by ultracentrifugation plus DGUC. The study^16^ also suggested, however, that EV protein content might differ more according to density rather than particle size. Hence, a future direction for bdEV studies might be explore the relationships between size and density more stringently, perhaps by using finer size-based fractionation such as asymmetric flow field-flow fractionation combined with density gradient separations.

We propose that our bdEV dataset can suggest important tools for biomarker discovery and mechanistic studies in neurological disease. For example, capturing bdEVs in peripheral samples can assist with monitoring neuropathological changes in the brain. Recently, L1CAM and NCAM ^1, 63, 64^ have been used to capture plasma neuron-derived EVs, and GLAST for astrocyte-derived EVs^65^. Our data may suggest additional CNS cell-enriched markers that could be used to capture or characterize bdEVs in the periphery. For example, neuron-specific markers detected here include enolase 2 (ENO2) and vesicle-associated membrane protein 2 (VAMP2). Also, many small RNA and protein components of bdEVs are involved in neuronal functions and neurodegenerative diseases. Of the miRNAs highly enriched in brain-derived EVs from both human and macaque samples (Figure 6G), miR-21 was reported to have neurotoxic effects in bdEVs in simian immunodeficiency virus (SIV)-induced central nervous system (CNS) disease^15^, while miR-7^66^, miR-125b^66^, and miR-222^67, 68^ were up-regulated in brain tissue^66^, CSF^68^ or plasma^67^ of Alzheimer’s disease (AD) patients. Based on protein findings, functional annotations and pathway analyses support contributions to neuronal functions. Tau protein is extensively used as a diagnostic tool for AD dementia^69^. TMEM30A interacts with the β carboxyl-terminal fragment of amyloid-β (Aβ) precursor protein in endosomes^70^ and is released from the blood-brain barrier^71^. Members of the 14-3-3 protein family and DJ-1 confirm previous findings in CSF derived EVs^72^. We expect that ongoing analyses of these data and future studies of bdEVs from different neurological diseases will yield further insights into mechanisms of neuropathology.

In summary, we characterized bdEVs from human, macaque, and mouse tissue harvested under different conditions, testing several variations on EV separation. We hope that this study will add to the growing understanding of the small RNome and proteome composition of bdEVs. We further trust that this study and the associated data will be used to further our knowledge of the regulatory roles of EVs in brain and to facilitate biomarker discovery for neurological diseases.

## Acknowledgments

This work was supported by grants from the US National Institutes of Health (DA040385, DA047807, AI144997, and MH118164, to KWW) and AG057430 (to VM and KWW); by the Michael J. Fox Foundation (to KWW); by UG3CA241694, supported by the NIH Common Fund, through the Office of Strategic Coordination/Office of the NIH Director; and by the National Center for Research Resources and the Office of Research Infrastructure Programs (ORIP) and the National Institutes of Health, grant number P40 OD013117. This work was also supported by the National Health and Medical Research Council of Australia, GNT1132604 (to AFH). We thank Suzanne Queen, Brandon Bullock, and Emily Mallick from the Johns Hopkins University School of Medicine for generously providing and processing the macaque brain tissues. We thank Connie Talbot, Bob Cole and Tatiana Boronina from the Johns Hopkins University School of Medicine for contributing to proteomics data analysis. We gratefully acknowledge the La Trobe University Comprehensive Proteomics Platform.

## Contributions of Authors

K.W.W, Y.H. and L.Z. conceived the idea. Y.H. performed most experiments and drafted and revised the manuscript; K.W.W. directed the project, obtained funding, supervised the experiments, and revised the manuscript; L.C and A.F.H performed small RNA libraries construction, small RNA sequence, mass spectrometry, and proteomics data analysis; A.T. performed small RNA sequencing bioinformatics analysis; V.M., J.C.T, and O.P. processed and provided the human brain samples; N.J.H. provided nanoparticle tracking analysis techniques; L.J.V developed the original protocol and helped with protocol modification. All authors read and approved the final manuscript.

---

**Figure S1.**
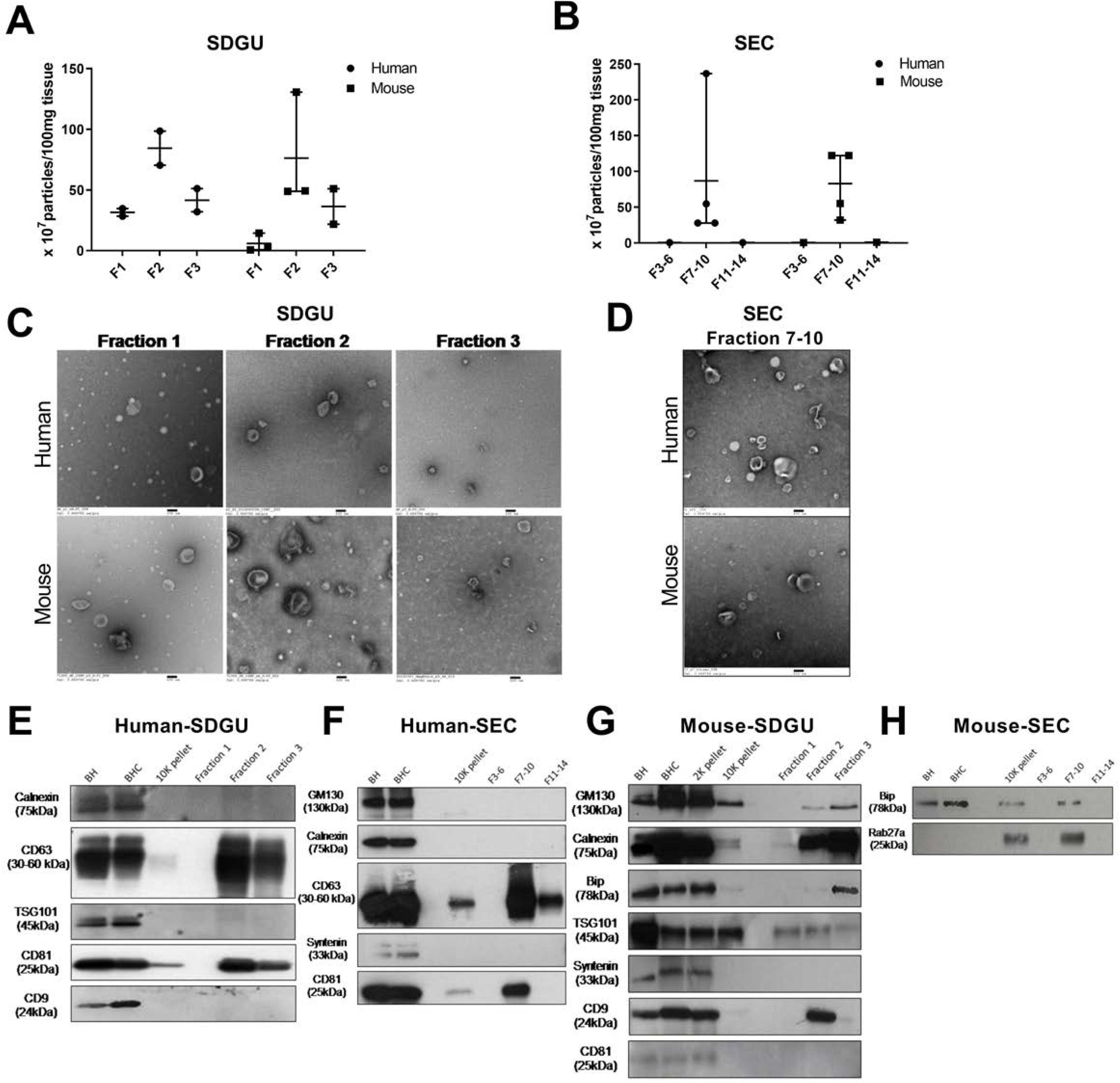
Comparison of bdEVs in different fractions of SDGU and SEC: human and mouse brain. (A) Particle concentration of SDGU fractions from human and mouse brain by NTA (Particle Metrix). (B) Particle concentration of SEC fractions from human and mouse brain (NTA). (A)-(B): Data are presented as the mean with range. (C) Three fractions from SDGU visualized by TEM (scale bar = 100 nm). (D) Fractions 7-10 from SEC (TEM, scale bar = 100 nm). TEM is representative of five images taken of each fraction from three independent brain tissue samples. (E) Western blot analysis of calnexin, CD63, TSG101, CD81, and CD9 of brain tissue and SDGU fractions (human) (n=1). (F) Western blot analysis of GM130, calnexin, CD63, syntenin, and CD81 of brain tissue and SEC fractions (human) (n=1). (G) Western blot analysis of GM130, calnexin, Bip, TSG101, syntenin, CD9, and CD81 of brain tissue and SDGU fractions (mouse) (n=1). (H) Western blot analysis of Bip and Rab27a of brain tissue and SEC fractions (mouse) (n=1).

**Figure S2.**
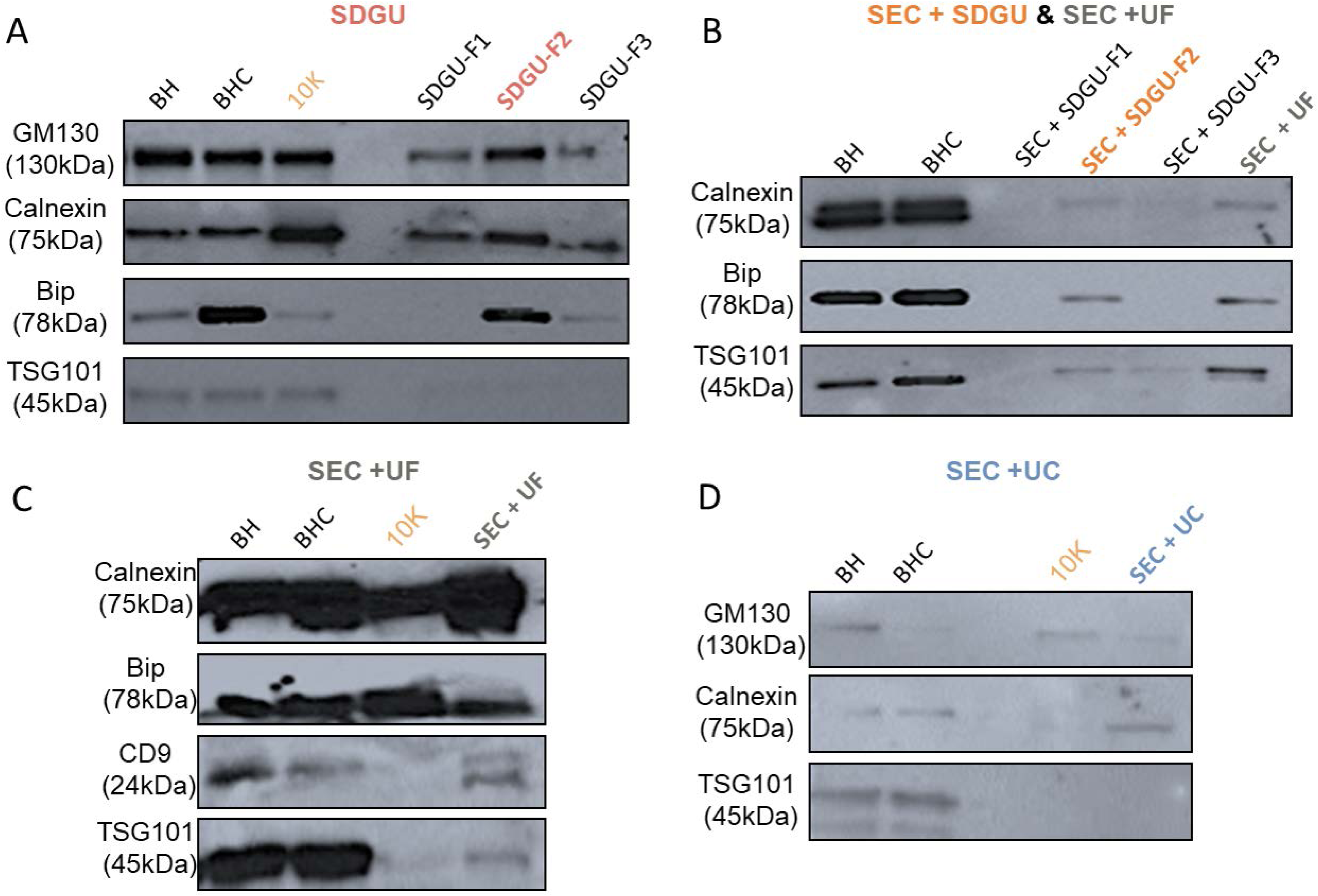
Western blot analysis: mouse brain tissue and EV preparations. In each of the following, tissue proteins were compared with EV fractions obtained as indicated. (A) SDGU fractions; immunoblotting for GM130, calnexin, Bip, and TSG101 (n=3). (B) SEC + SDGU and SEC + UF fractions; immunoblotting for calnexin, Bip, and TSG101 (n=1). (C) SEC + UF method fractions; immunoblotting for calnexin, Bip, CD9, and TSG101 (n=2). (D) SEC + UC; immunoblotting for GM130, calnexin, and TSG101 (n=1).

**Figure S3.**
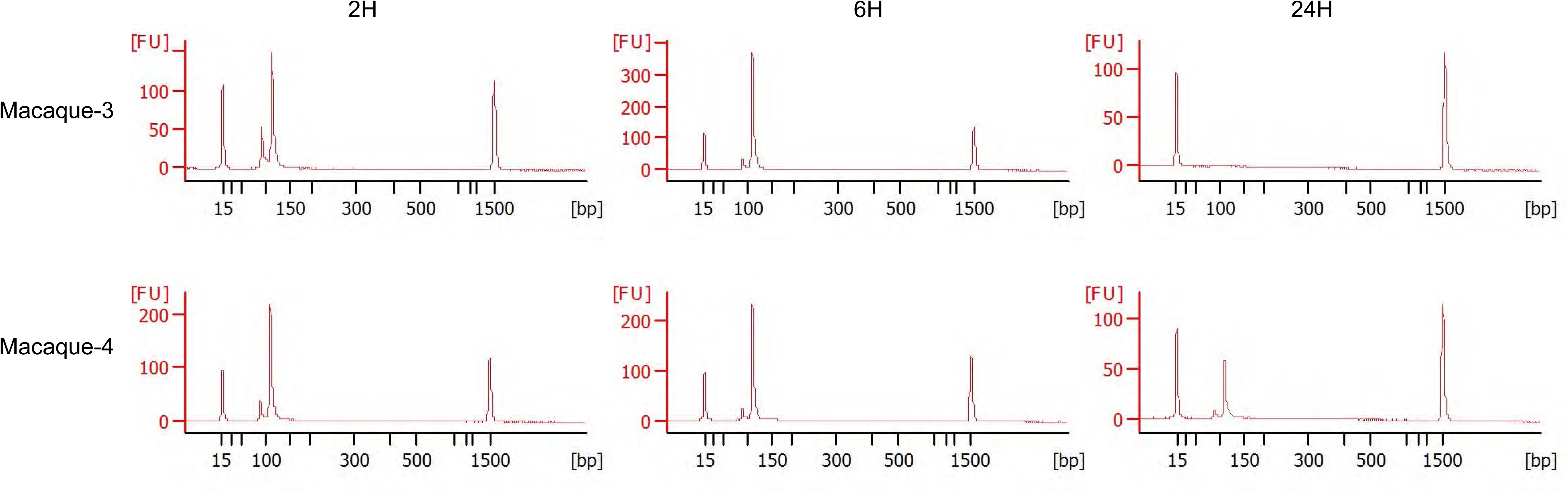
Bioanalyzer analysis of bdEV small RNA libraries. Size distribution of bdEV libraries from 2H, 6H, and 24H postmortem interval (PMI) macaque brains (n=2), measured by Bioanalyzer.

**Figure S4.**
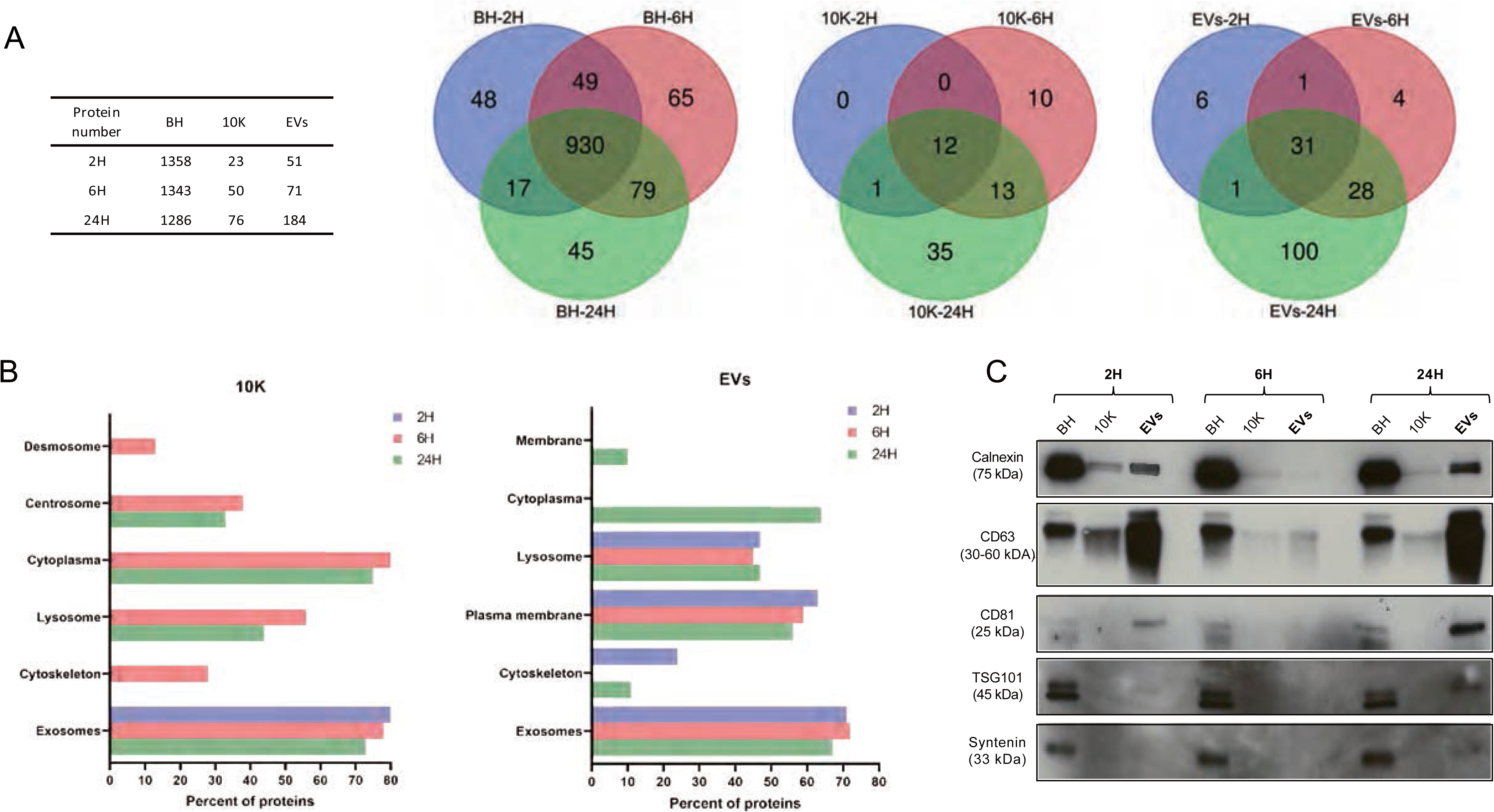
Effect of postmortem interval on bdEV protein contents. (A) Venn diagram of identified protein number of BH, 10K, and EVs at 2H, 6H, and 24H PMI in either two macaques. (B) Enrichment of cellular compartments of proteins identified in 10K and EVs at 2H, 6H, and 24H PMI. Gene ontology (GO) analysis was performed with Funrich. GO terms with FDR-corrected p-value < 0.05 are shown). (C) Western blot analysis of calnexin, CD63, CD81, TSG101, and syntenin associated with BH and EVs of macaque brain tissue at different PMI. Blots are representative of two independent tissue EV separations.

**Figure S5.**
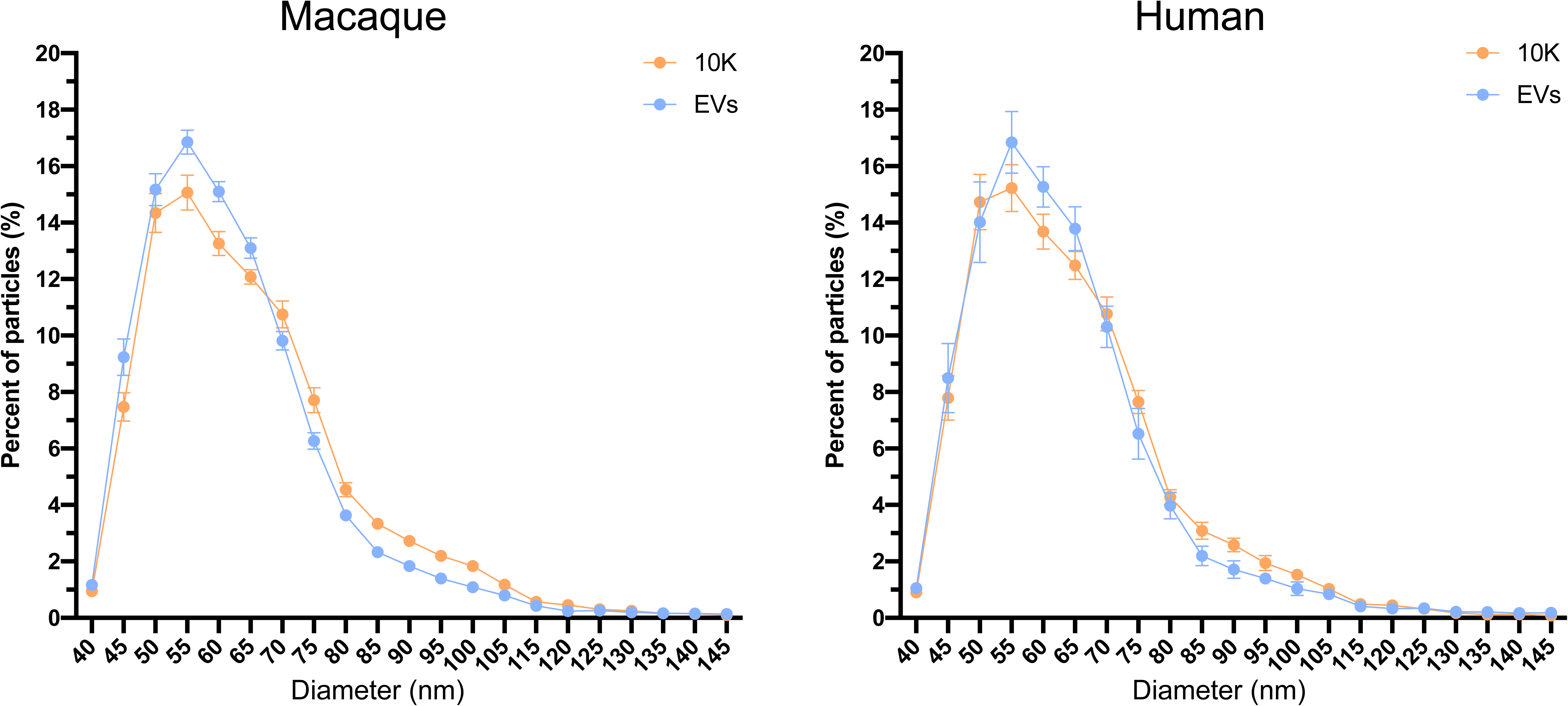
Size distribution of bdEVs from macaque and human brain as determined by nanoFCM. The size distribution for bdEV separated by SEC+UC from human and macaque was measured by nanoFCM flow nano-Analyzer (NanoFCM Co.). Data are presented as mean with standard deviation (n=2 for macaque, n=7 for human).

**Figure S6.**
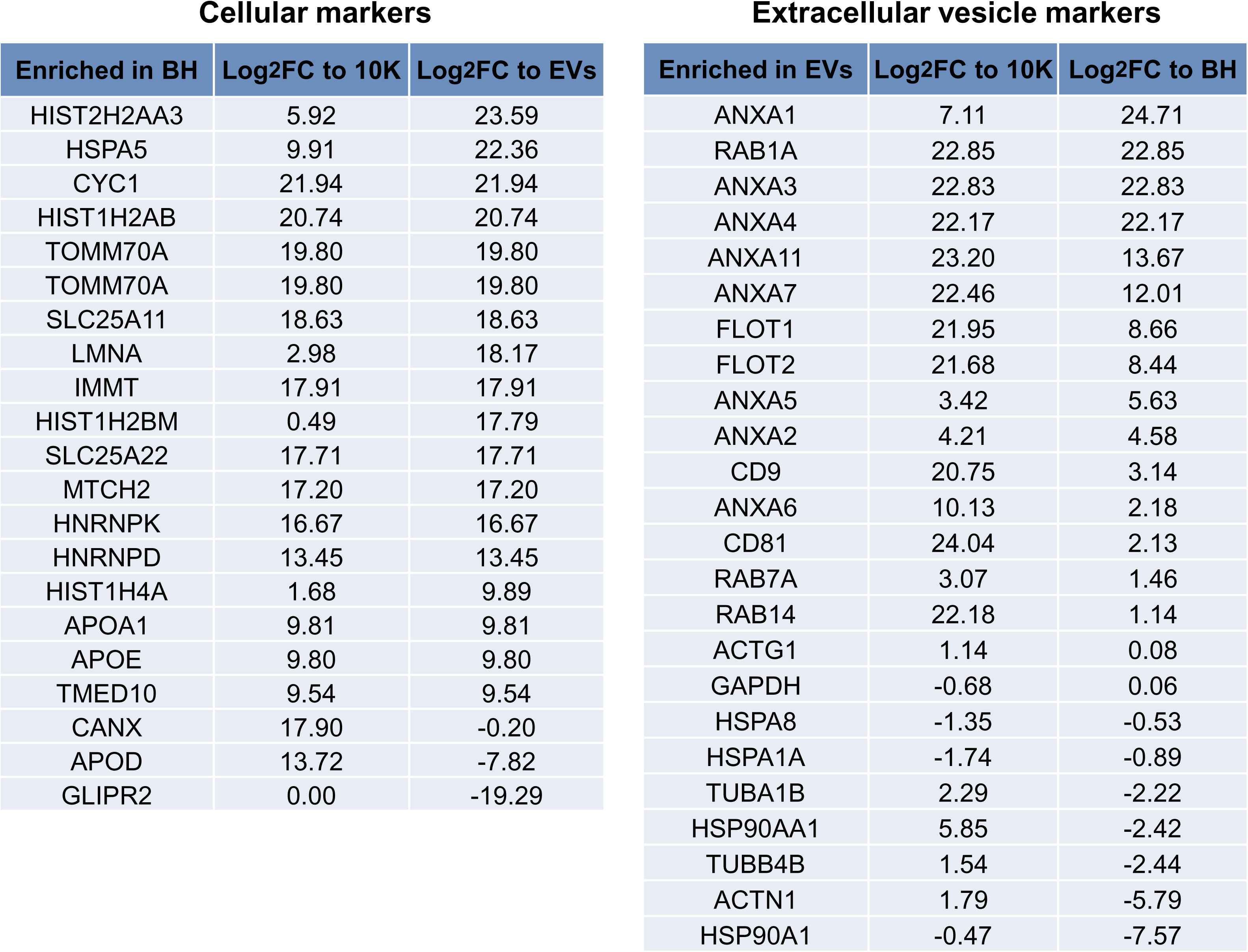
Enrichment of cellular and extracellular vesicle marker proteins in human brain tissue and EVs. Log_2_ fold-change (Log_2_FC) of cellular proteins enriched in BH compared with 10K and EVs (left panel) and extracellular proteins enriched in EVs compared with 10K and BH (right panel) (n=7).

